# 3D MAPs discovers the morphological sequence chondrocytes undergo in the growth plate and the regulatory role of GDF5 in this process

**DOI:** 10.1101/2020.07.28.225409

**Authors:** Sarah Rubin, Ankit Agrawal, Johannes Stegmaier, Jonathan Svorai, Yoseph Addadi, Paul Villoutreix, Tomer Stern, Elazar Zelzer

## Abstract

The activity of the epiphyseal growth plates, which drive longitudinal growth of long bones, is dependent on the ability of chondrocytes to change their shape and size extensively as they differentiate. However, organ size, extracellular matrix density and cell number have hindered the study of chondrocyte morphology. Here, we describe a new pipeline called 3D Morphometric Analysis for Phenotypic significance (3D MAPs), which overcomes these obstacles. By using 3D MAPs, we have created an image database of hundreds of thousands of cells from orthologous long bones. Analysis of this database revealed the growth strategies that chondrocytes use during differentiation. We found that chondrocytes employed both allometric and isometric growth, and that allometric growth is achieved by changes either in volume or surface area along a specific cell axis in a zone-specific manner. Additionally, we discovered a new organization of chondrocytes within the growth plate, where cells are orientated such that their longest axis always aligns with the dorsal-ventral axis of the bone. To demonstrate the ability of 3D MAPs to explore mechanisms of growth plate regulation, we studied the abnormally short tibiae of *Gdf5-*null mice. 3D MAPs identified aberrant cellular growth behaviors which resulted in a 3-fold reduction in volumetric cell growth, as well as affected cell morphology and orientation, highlighting GDF5 as a new regulator of growth plate activity. Overall, our findings provide new insight into the morphological sequence that chondrocytes undergo during differentiation and highlight the ability of 3D MAPs to uncover molecular and cellular mechanisms regulating this process. More broadly, this work provides a new framework for studying growth plate biology.

## Introduction

The linkage between cell morphology and organization and the architecture of a tissue is well-recognized. For example, during early fly development, cell rearrangements such as intercalation and rosette formation are the main driving force of germ-band extension, while apical cell constriction drives mesoderm invagination (Irvine and Wieschaus 1994, Hogan 1999, Lecuit and Lenne 2007, Martin, Kaschube et al. 2009, Paluch and Heisenberg 2009, Kim and Davidson 2011). Thus far, most of these morphogenetic studies have focused on two-dimensional epithelial sheets or early embryogenesis (Ingber, L. et al. 1994, Rauzi and Lenne 2011, Stegmaier, Amat et al. 2016, Heer and Martin 2017, Sutherland, Keller et al. 2019, Diaz-de-la-Loza, Loker et al. 2020). Studying these cellular behaviors in large three dimensional (3D) tissues is challenging, as the data requires the development of a methodology for time-efficient 3D imaging, registration, segmentation, and quantitative analysis of cells. An additional challenge is investigating these processes in mice, where tissues and organs can contain hundreds of thousands of cells, which forms multiple data scales. Focusing on the tissue level results in loss of subcellular resolution, while focusing on smaller regions at high resolution results in loss of their relationship to 3D tissue morphology. Recent advances in tissue clearing combined with light-sheet fluorescence microscopy has allowed for high-resolution mapping of fine tissue architectures in intact 3D samples (Yang, Treweek et al. 2014, Calve, Ready et al. 2015, Treweek, Chan et al. 2015, Greenbaum, Chan et al. 2017). Yet, despite the advances in whole-organ imaging, there is still a lack of tools with which to accurately extract the 3D morphology of cells and identify morphological patterns in large 3D datasets in a time-efficient manner.

The growth plate is an interesting model for studying 3D cell behavior as a driver of tissue morphogenesis. Growth plates are cartilaginous tissues located at either end of developing bones, whose function is to drive bone elongation (Cancedda, Cancedda et al. 1995, Noonan, Hunziker et al. 1998, St-Jacques, M. et al. 1999, Kronenberg 2003, Wilsman, Bernardini et al. 2008).

Typically, the growth plate is divided into four zones along the proximal-distal (P-D) axis of the bone. Most extreme is the resting zone (RZ), then the proliferative zone (PZ) followed by prehypertrophic (PHZ) and hypertrophic zones (HZ). This arrangement reflects a series of differentiation states that are marked by unique cell morphologies, extracellular matrix (ECM) properties and gene expression profiles (Cancedda, Cancedda et al. 1995, Abad, Meyers et al. 2002, Kronenberg 2003, Mackie, Ahmed et al. 2008, Cooper, Oh et al. 2013, Kozhemyakina, Lassar et al. 2015).

Several studies have aimed to decipher cell behaviors affecting growth plate structure and function. One example of such behaviors is the massive swelling that occurs when cells undergo hypertrophy. This 9-fold increase in cell volume drives bone elongation (Hunziker, Schenk et al. 1987, Hunziker and Schenk 1989, Breur, Vanenkevort et al. 1991, Wilsman, Farnum et al. 1996, Breur, Lapierre et al. 1997, Wilsman, Bernardini et al. 2008, Cooper, Oh et al. 2013, Li, Trivedi et al. 2015, Cooper 2019). Another cell behavior linked to growth plate function is column formation. When chondrocytes enter the proliferative zone, they flatten and arrange themselves into columns (Dodds 1930, Aszodi, Hunziker et al. 2003, Li and Dudley 2009, Li, Li et al. 2017). The orientation of the short axis of these flattened cells becomes parallel to the P-D axis of the bone (Li and Dudley 2009, Romereim, Conoan et al. 2014). The significance of this process was demonstrated in mice lacking key components of the planar cell polarity (PCP) pathway or Beta1 integrins, which displayed short and misshapen bones due to impaired column formation (Aszodi, Hunziker et al. 2003, Li and Dudley 2009, Gao, Song et al. 2011, Shwartz, Farkas et al. 2012, Li, Li et al. 2017). Together, these works demonstrate that cell shape and orientation are essential drivers of growth plate structure and function.

Notwithstanding these important discoveries, comprehensive understanding of growth strategies of differentiating chondrocytes and the relation between their morphology and growth plate structure and function is still lacking. One reason for that is that due to high density of cells and ECM, imaging the growth plate in its entirety has been a major challenge. Additionally, there is a lack of segmentation tools to accurately extract entire cells for analysis. To overcome these obstacles, we have developed a modular imaging and analysis pipeline called 3D MAPs, which enables to explore large datasets of hundreds of thousands of cells, accurately characterizing their morphology while preserving their tissue-level position. Using 3D MAPs, we discovered new cellular behaviors and patterns that contribute to growth plate structure and activity. Moreover, utilizing this pipeline we identified abnormal cellular behaviors of chondrocytes in growth plates of *Gdf5*-null mice, exposing a new role for this gene in skeletogenesis and concomitantly underscoring the ability of 3D MAPs to identify new molecular components in growth plate biology.

## Results

### 3D MAPs: Morphometric Analysis to test for Phenotypic significance

There are likely key aspects of cell morphology that are important for growth plate function, but there is currently no tool suitable for mining this type of data. To that end, we developed a pipeline called 3D Morphometric Analysis to test for Phenotypic significance (3D MAPs). The major steps of 3D MAPs are clearing, imaging, segmentation, and 3D morphometric analysis. To overcome the challenge of imaging a large and dense tissue, we performed tissue clearing, which enabled us to image the entire growth plate by light-sheet microscopy (Figure 1A and Bi,ii). To accurately segment entire cells, we designed a new segmentation pipeline in an XML pipeline wrapper for the Insight Toolkit (XPIWIT) (Bartschat, Hübner et al. 2015), which automatically segments cells and their nuclei (Figure 1Biii). Finally, to correlate between cell shape and its position in the growth plate, we extracted various morphological features (Supplementary Table 1) and then projected them onto the cell location in the growth plate, generating 3D morphology maps (Figure 1Biv). We could then visually identify potentially important patterns in morphology and analyze these features quantitatively and comparatively across space.

**Figure 1.**
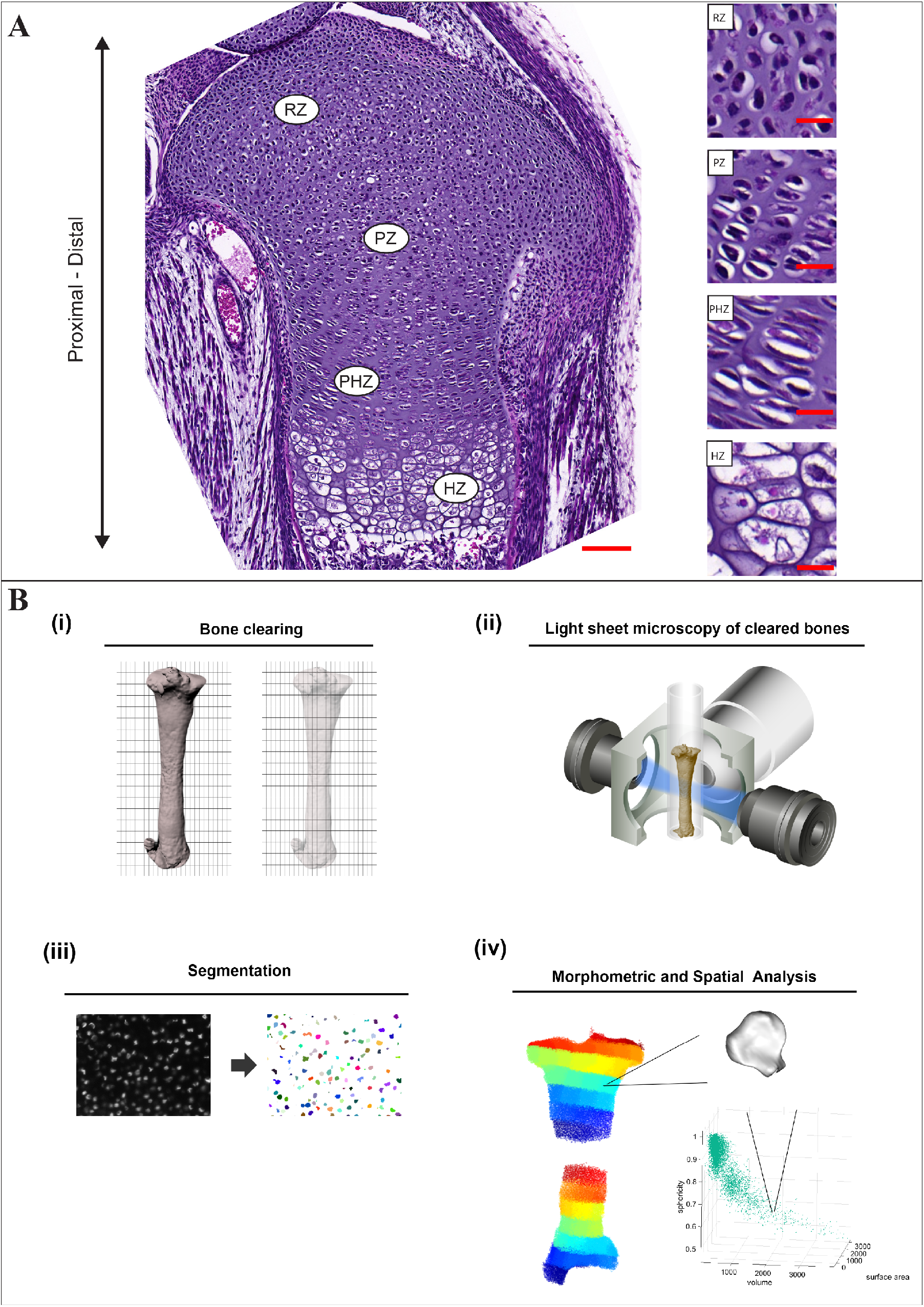
3D MAPs: Morphometric Analysis to test for Phenotypic significance. (**A**) 7 μm-thick paraffin sections of an E18.5 proximal tibial growth plate stained with H&E show the four zones of the growth plate along the proximal-distal axis of the bone. Scale bar: 100 μm. The magnified images on the right show the morphological changes of chondrocytes during their differentiation sequence, including flattening and column formation of proliferative zone (PZ) chondrocytes, enlargement of column cells in the prehypertrophic zone (PHZ) and increase in size of terminally differentiated chondrocytes in the hypertrophic zone (HZ). Scale bars: 20 μm. (**B**) An overview of 3D MAPs pipeline. (i) The bone is dissected and cleared by PACT-deCAL, and nuclei are fluorescently labeled. (ii) The cleared bone is embedded in a glass capillary and proximal and distal growth plates are imaged at high resolution with a Z.1 light sheet microscope. (iii) Each image stack undergoes nuclear and cellular segmentation. (iv) The segmented cells and nuclei are subjected to morphometric analyses and are projected back onto their anatomical position in the bone.

### Data acquisition and segmentation

For tissue clearing, we used the PACT-deCAL technique (Yang, Treweek et al. 2014, Treweek, Chan et al. 2015), which was found to allow nuclear labeling while preserving endogenous tissue fluorescence. Next, E16.5 nucleus- and cell membrane-labeled growth plates from tibias and ulnas were imaged by light-sheet fluorescence microscopy (LSFM) (Supplementary Figure 1A). Due to its high speed, LSFM proved to be the superior choice over other laser-based techniques, such as confocal and two-photon microscopy (data not shown). To reduce the file size and expedite image processing, acquired z stacks included only regions with growth plate nuclei and cells and images were downsampled in XY to a resolution lower than the cell membrane thickness prior to segmentation.

The next challenge was to conduct accurate and time-efficient 3D segmentation of nuclei and cells from the light-sheet images. For that, we combined two open-source platforms to perform semi-automatic segmentations (SAS) (Supplementary Figure 1B). Nuclei and cells from surrounding tissues were manually excluded by masking using Microview 2.1.2 (GE Healthcare). Next, the cleaned raw images were automatically segmented using specially designed pipelines for the nuclei and cells in XPIWIT software (Stegmaier, Otte et al. 2014, Bartschat, Hübner et al. 2015). Then, nuclei and cells were filtered based on volume, and cells were analyzed only if they had a matching nucleus smaller in volume than the cell itself. On average, this segmentation stage resulted in the correct identification of 50% of nuclei and 25% of cells from all growth plate samples; for example, 124,400 nuclei and 62,200 cells in the proximal tibial growth plate (Supplementary Figure 1C).

To verify the accuracy of segmentations, we performed a benchmark analysis comparing manual and automatic segmentations from two different imaging modalities, namely confocal and light-sheet microscopy. To establish the “ground truth” (GT), nuclei and cells from the four growth plate zones were manually segmented from confocal images in Microview. Volume and surface area were extracted, and corresponding histograms were compared between manual and semi-automatic segmentations using chi-square distance test. Results showed that nucleus and cell segmentations by SAS were indistinguishable from manual GT segmentations (p > 0.05), demonstrating the validity of our measurements (Supplementary Figure 1D).

### 3D MAPs can identify morphogenetic behaviors in the growth plate

Having segmented successfully cells from each growth plate, we proceeded to characterize various features of cell morphology (listed in Supplementary Table 1) in three growth plates from forelimb and hindlimb bones, namely proximal tibia (PT), distal tibia (DT), and distal ulna (DU). The proximal ulnar growth plate was excluded from our analysis because it has a very complex morphology. First, we analyzed well-known properties of cells, such as volume, surface area and density. In agreement with previous reports on chondrocyte hypertrophy (Cooper, Oh et al. 2013), cells increased their volume on average 9-fold from the resting to hypertrophic zone (Figure 2A). Our analysis further revealed that on average, cells in the proliferative zone grew 20% larger than resting zone cells, prehypertrophic cells grew 74% larger than proliferative zone cells and hypertrophic cells grew 50% larger than prehypertrophic cells. Additionally, we found that cells increased their surface area 5-fold on average as they underwent differentiation, increasing by 20% from the resting to proliferative zone, 66% from the proliferative to prehypertrophic zone, and 36% from the prehypertrophic to hypertrophic zone. Cell density was more variable between the three growth plates, but it decreased on average 5-fold during differentiation (12-fold at most). In the distal tibia for example, cell density increased by 7% from the resting to proliferative zone, and then decreased by 60% from the proliferative to prehypertrophic zone, and again by 37% from the prehypertrophic to hypertrophic zone (Figure 2A).

**Figure 2.**
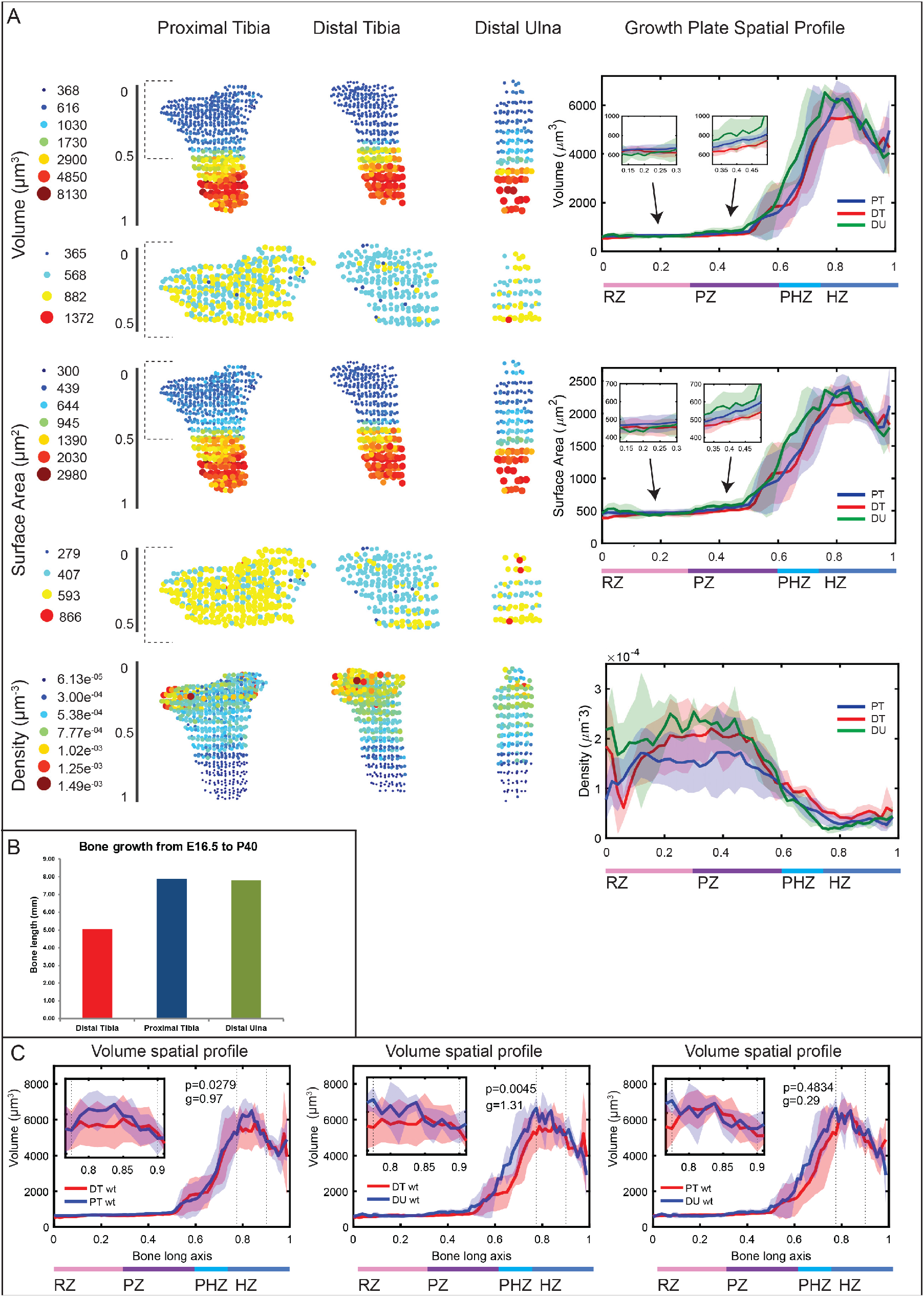
3D MAPs can identify morphogenetic behaviors in the growth plate. Morphology and spatial analysis were performed for each segmented cell and each feature was projected back to the anatomical position of that cell, creating a 3D growth plate map. Each circle represents the mean value within a 75 μm cube. All 3D maps are oriented such that the RZ is at the top and the HZ is at the bottom. Additionally, all growth plates were registered to one another, and a spatial profile was created showing how each feature changes along the differentiation axis of each growth plate. (**A**) Cell volume, surface area, and density were computed for the growth plates of the proximal tibia (PT), distal tibia (DT), and distal ulna (DU). Dashed grey brackets demarcate zoomed-in area of the first half (0 - 0.5) of each growth plate, highlighting variations in volume and surface area among the different growth plates. Spatial profiles, which were plotted for each feature, show that in all three growth plates, cell volume and surface area increased during chondrocyte differentiation, while cell density decreased. Shaded region shows standard deviation between samples (PT and DT, n = 5; DU, n = 3). Insets in graphs show zoomed-in regions from the RZ and PZ to highlight the volume and surface area increase. (**B**) Bar graph shows mean contribution of PT (blue), DT (red) and DU (green) growth plates to total bone growth from E16.5 to P40 (n = 3). The PT and DU growth plates were more active than the DT growth plate. (**C**) The largest 10% of HZ cell volumes (region between dashed lines) were compared by Student’s *t*-test between pairs of growth plates. Correlating with the growth plate activity, HZ cell volumes were significantly different between DT and PT (p = 0.0279, g = 0.97) and between DT and DU growth plates (p = 0.0045, g = 1.31), but not significantly different between PT and DU growth plates (p = 0.4834, g = 0.29).

There is a well-established positive correlation between hypertrophic cell volume and growth potential of different growth plates (Breur, Vanenkevort et al. 1991, Cooper, Oh et al. 2013, Cooper 2019). To verify that in our datasets, we calculated the growth of the PT, DT, and DU growth plates from E16.5 to postnatal day (P) 40 using previously published data (Stern, Aviram et al. 2015) (Figure 2B) and compared them to the top 10% hypertrophic cell volumes from these growth plates at E16.5 (Figure 2C). Consistent with previous studies, hypertrophic cell volumes differed between growth plates and correlated with their activity. The DT growth plate grew less than both PT and DU, and also had significantly smaller cell volumes in the hypertrophic zone. PT and DU growth plates grew at the same rate and displayed similar hypertrophic cell volumes. Together, these findings confirm the ability of 3D MAPs to identify morphogenetic behaviors of cells in the growth plate.

### Allometric and isometric growth behaviors define chondrocyte shape throughout differentiation

Cell growth can be either isometric or allometric. During isometric growth, volume scales to the two-third power compared to the surface area (McMahon, Bonner et al. 1983, Okie 2013). To determine whether chondrocyte growth during differentiation is isometric or allometric, we plotted the average volume^2/3^ to surface area (Vol^2/3^/SA) ratio along the growth plate and used linear regression to characterize the growth as isometric (slope = 0) or allometric (slope ≠ 0). Interestingly, we identified four main growth strategies along the growth plate (Figure 3A and 3A’). In the proximal tibia and distal ulna, resting zone cells started to grow allometrically, driven by volume increase. As they differentiated to the proliferative zone, volume increase slowed down and allometry was driven by an increase in surface area (slope < 0). In the prehypertrophic zone, cells changed their growth behavior again and allometry was driven by volume (slope > 0). Finally, hypertrophic cells changed their growth strategy to be isometric (slope = 0). The distal tibia growth plate displayed the same trends except in the resting zone, where cells grew allometrically driven by surface area. The consistency of these findings across growth plates suggests the generality of this finding, whereas the difference in the distal tibia resting zone may reflect its different growth rate, as compared to the two other growth plates.

**Figure 3.**
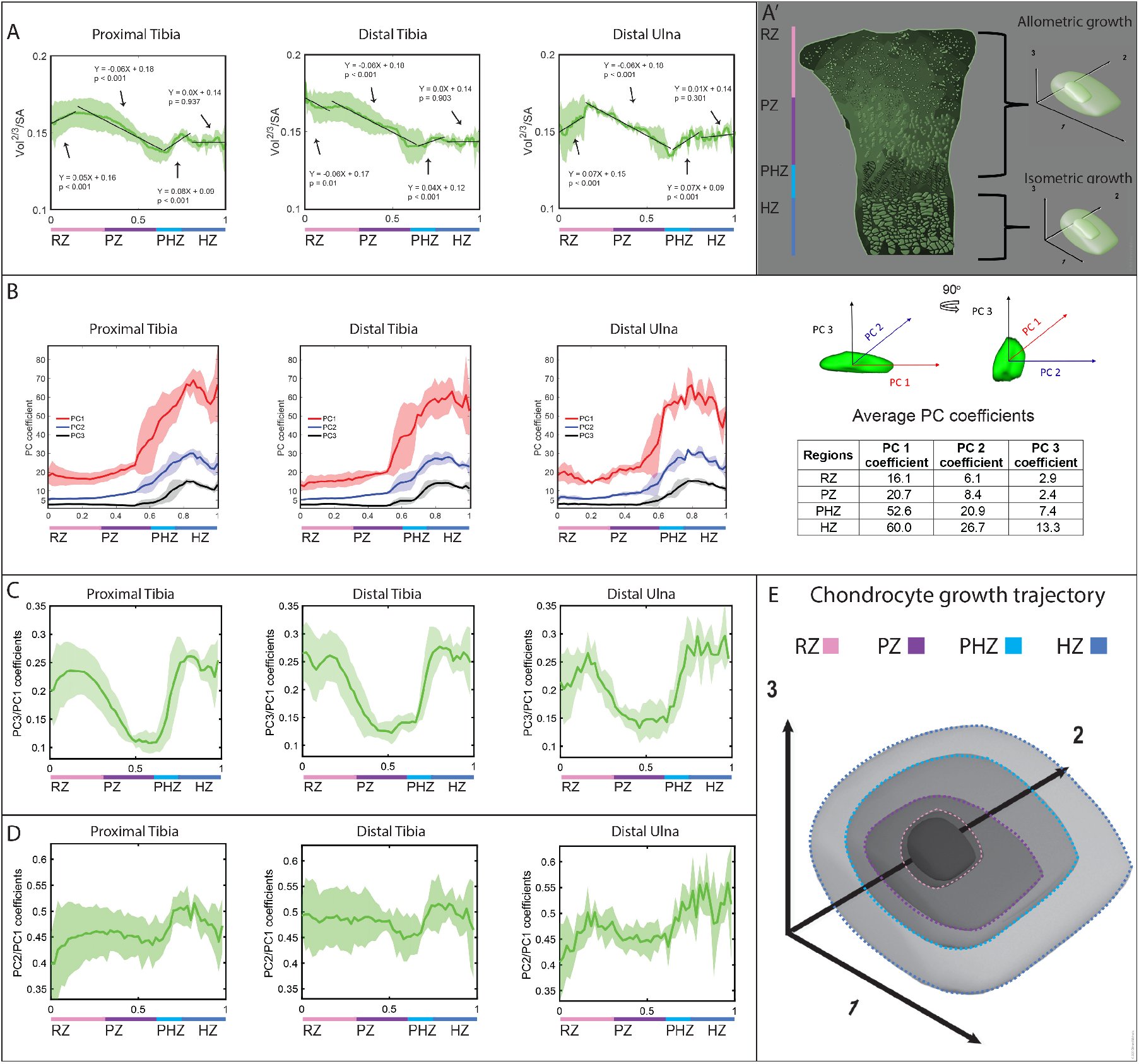
Allometric and isometric growth behaviors define chondrocyte shape throughout differentiation. **(A)** The Vol^2/3^/SA ratio was plotted as a spatial profile along the differentiation axis and regression lines were fit for each change in growth behavior to determine if the growth was isometric (slope = 0) or allometric (slope ≠ 0). Slopes > 0 represent volume-dependent allometric growth, whereas slopes < 0 represent surface area-dependent allometric growth. In all three growth plates, four identified growth behaviors correlated with the differentiation state. In the PT and DU, RZ cells grew by volume-dependent allometric growth (PT slope = 0.05 p = 5.92e^−09^ R^2^ = 0.93; DU slope = 0.07 p = 7.37e^−04^ R^2^ = 0.6), while in the DT, RZ cells grew by surface area-dependent allometric growth (DT slope =−0.06 p = 1.06e^−02^ R^2^ = 0.41). The rest of the growth behaviors were conserved among all three growth plates. In the PZ, cells grew by surface area-dependent allometric growth (PT slope = −0.06 p = 2.89e^−31^ R^2^ = 0.95; DT slope = −0.06 p = 5.71e^−33^ R^2^ = 0.95; DU slope = −0.06 p = 2.06e^−34^ R^2^ = 0.96), in the PHZ by volume-dependent allometric growth (PT slope = 0.08 p = 8.09e^−5^ R^2^ = 0.68; DT slope = 0.04 p = 3.11e^−4^ R^2^ = 0.62; DU slope = 0.07 p = 8.64e^−4^ R^2^ = 0.56), and in the HZ cells grew by isometric growth (PT slope = 0 p = 0.937 R^2^ < 0.01; DT slope = 0 p = 0.903 R^2^ < 0.01; DU slope = 0.01p = 0.301 R^2^ = 0.06). (**A’**) Scheme illustrating that chondrocytes change their shape as they grow allometrically in the RZ, PZ, and PHZ, but maintain their shape while growing isometrically in the HZ. (**B-D**) Principal component analysis (PCA). (**B**): PC1 (red arrow) of the object represents the longest cell axis, PC2 (blue arrow) the second longest axis, and PC3 (black arrow) the shortest axis. Spatial profiles of PC 1, 2, and 3 show that they are significantly larger than one another and they are conserved among the three growth plates. The table shows the mean PC coefficients across all three growth plates per zone. (**C**): In all three growth plates, as cells differentiated from the RZ to the PZ, they decreased their PC3/PC1 ratio by half and then returned to the same ratio when they differentiated to the PHZ and HZ. (**D**): In the PT and DU, cells increased their PC2/PC1 ratio in the RZ and slightly decreased their ratio in the DT RZ. In all three growth plates, the PC2/PC1 ratio was constant throughout the PZ, and increased as cells differentiate to the PHZ. The ratio at the end of the HZ is the same as in the RZ. (**E**) Scheme illustrating how each cell axis changes during growth and differentiation in the growth plate. (Shaded region shows standard deviation between samples (PT and DT, n = 5; DU, n = 3). p-values were calculated by Student’s *t*-test between slope = 0 and slope of fit line. Abbreviations: PT = proximal tibia; DT = distal tibia; DU = distal ulna; RZ = resting zone; PZ = proliferative zone; PHZ = prehypertrophic zone; HZ = hypertrophic zone.

To connect between the different growth strategies and changes in cell morphology, we followed the changes in ratios between the three cell axes during differentiation. To extract cell axes, we performed principal component analysis (PCA) on the masked region of the segmented cell (Figure 3B), where PC3 describes the shortest cell axis, PC2 describes the second longest and PC1 describes the longest axis. This analysis identified large variances of at least 2-fold between all three principal components, suggesting that their assignments can be used to represent the three cell axes (Figure 3B). To capture morphological changes during differentiation, the PC coefficients were plotted as ratios along the growth plate (PC3/PC1, PC2/PC1, and PC3/PC2) (Figure 3C, 3D and Supplementary Figure 2). As seen in Figure 3C, comparing the shortest to longest cell axes (PC3/PC1) we found that cells in the resting zone start at a ratio of 0.25 and then decrease by 50% in the proliferative zone. At the same time, the ratio of the second longest to longest cell axis (PC2/PC1) decreases slightly in this transition (Figure 3D and 3E). Since this same region grows allometrically by surface area increase, we can deduce that cells flatten in this region by reducing their shortest axis (PC3) (Figure 3B and 3E) while increasing their surface area along their longest axis (PC1) and second longest axis (PC2). As cells continued to differentiate towards the prehypertrophic zone, the PC3/PC1 ratio as well as PC2/PC1 ratio started to increase. This means that cells grow allometrically in this region by specifically increasing volume along their shortest (PC3) and second longest (PC2) axes (Figure 3E). Once cells reached the hypertrophic zone, they had already reached their final shape and they swelled through isometric growth (Figure 3E).

Altogether, we found that both allometric and isometric growth strategies are employed by chondrocytes during differentiation. Remarkably, allometric chondrocyte growth is achieved by differentially directing changes in volume or surface area to a specific cell axis. These multiple levels of hierarchy suggest that chondrocyte growth during differentiation must be a tightly regulated process.

### Dorsal-ventral polarization of longest cell axis in the growth plate

Cell orientation in the growth plate is important, especially in the proliferative zone, where cells orient their shortest axis parallel to the long axis of the bone to form columns, thereby facilitating bone elongation (Williams, Wr. et al. 2007, de Andrea, M et al. 2010, Kuss, Kraft et al. 2014, Romereim, Conoan et al. 2014). Previous studies also showed that the cell division occurs perpendicular to the axis of growth. However, these analyses were performed in 2D, so nothing is known about the orientation of the third cell axis and how this relates to what was previously described. To study in 3D the orientations of differentiating chondrocytes, we registered the growth plates and quantified the orientation of the three orthogonal axes of each cell using PCA (Figure 4A).

**Figure 4.**
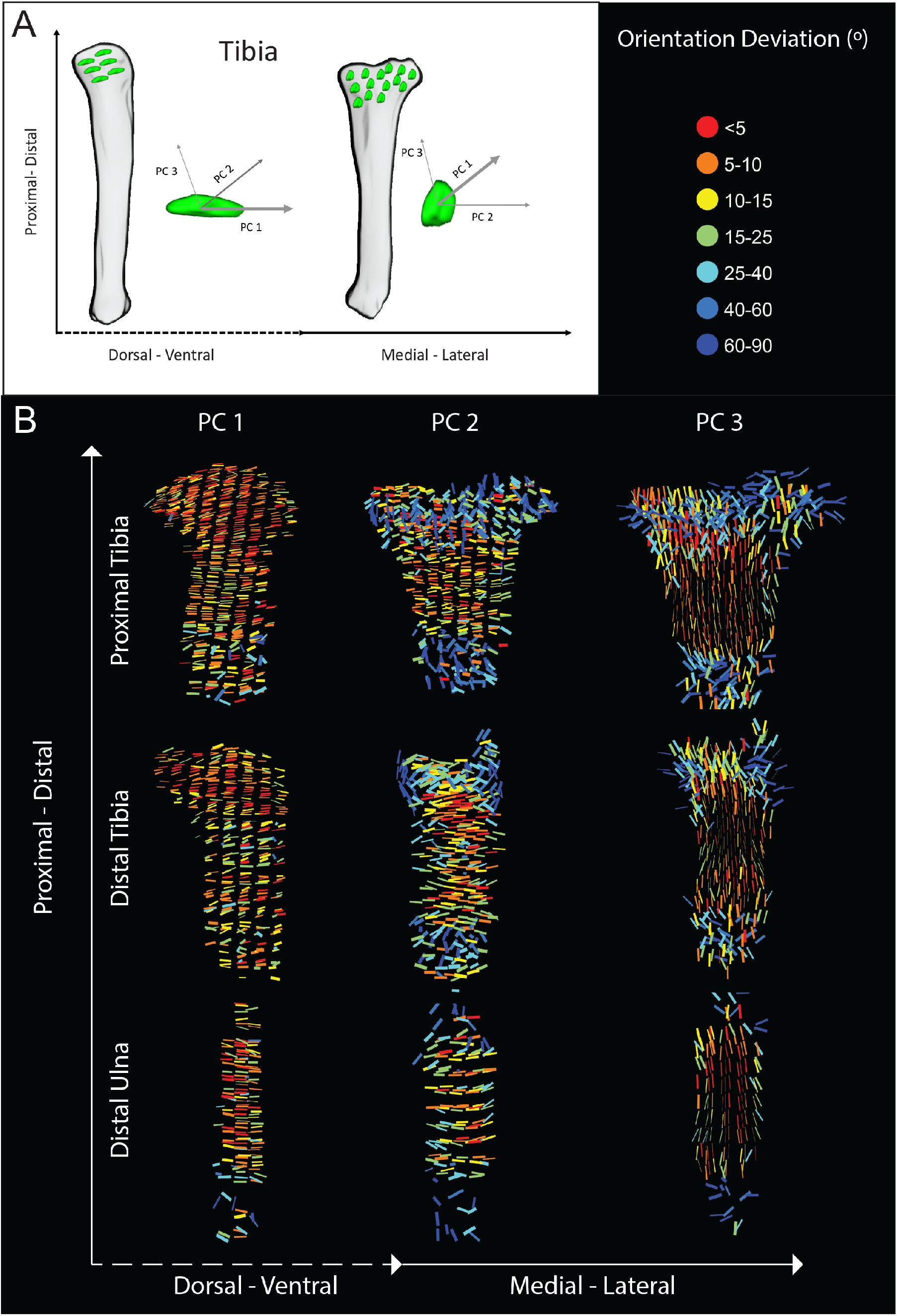
Dorsal-ventral polarization of longest cell axis in the growth plate. Cell orientation was calculated by extracting the three orthogonal axes of each cell from the segmented object using principle component analysis (PCA). PC1 (thickest arrow) of the object represents the longest axis, PC2 (thinner arrow) the second longest axis, and PC3 (thinnest arrow) the shortest axis. The schematic drawing of the tibia in (**A**) shows that bone alignment determines which orientation axes can be viewed in 2D. (**B**) Cell orientation along the three axes (PC 1, 2, and 3) was plotted for all three growth plates (proximal tibia, distal tibia, and distal ulna). In all cases, the growth plates were placed such that the RZ pointed upward and the HZ pointed downward. The colormap is normalized to each sample, representing the deviation in degrees from the average orientation of each growth plate. All three growth plates exhibited similar cell orientation behaviors. PC1, the longest cell axis, aligned to the dorsal-ventral axis of the bone. The fact that most lines are red means that in each growth plate there was a strong preference to one direction, with some variability occurring in the HZ. PC2, the second longest axis, was highly variable in the RZ and HZ but aligned with high similarity to the medial-lateral axis of the bone. PC3, the shortest axis, was also highly variable in the RZ and HZ, but aligned with high similarity towards the proximal-distal axis in the PZ. Abbreviations: RZ = resting zone; PZ = proliferative zone; PHZ = prehypertrophic zone; HZ = hypertrophic zone.

To identify deviations in orientation, color maps were normalized to represent the angle between a cell's vector and the mean vector of all cells in the growth plate. In agreement with previous studies (Li and Dudley 2009, de Andrea, M et al. 2010, Romereim, Conoan et al. 2014), we found that proliferative zone cells align their shortest axis (PC3) along the proximal-distal axis of the bone (Figure 4B) with very little variation (indicated by the thin lines). In the resting and hypertrophic zones, we observed high variability in PC3 orientation (indicated by thick lines) and overall disorder. Prior works showed that the direction of cell intercalation in the proliferative zone is along the medial-lateral axis (Li and Dudley 2009, Prein, Warmbold et al. 2016, Li, Li et al. 2017). Interestingly, we found that this plane represents the second longest axis of the cell (PC2). Its profile was similar to PC3, where the proliferative zone was highly ordered with its axis orienting along the medial-lateral axis with low variability, but with increased variability in orientation in the resting and hypertrophic zones (Figure 4B).

Finally, we calculated the orientation of the longest cell axis (PC1), which arguably has the most potential to influence growth plate structure. Strikingly, we found that throughout the growth plate, the cells oriented PC1 towards the dorsal-ventral axis, with some variability occurring in the resting and hypertrophic zones (Figure 4B). Notably, the same patterns for all three orientation axes were conserved in the three analyzed growth plates. Altogether, by analyzing 3D cell orientation we discovered a new cell behavior in which the longest cell axis points towards the dorsal-ventral axis in all zones and in different growth plates, suggesting a biological need for this behavior.

### *Gdf5* regulates chondrocyte growth and morphology during differentiation

To demonstrate the sensitivity of 3D MAPs to identify abnormalities in cell growth and morphology and to identify new regulators of this process, we sought to study growth plates of mutants for genes whose loss of function led to abnormal skeletal development, yet their involvement in growth plate biology was unknown. A search of the literature revealed that loss of *Gdf5*, a well-known regulator of joint formation (Storm and Kingsley 1996, Settle, Rountree et al. 2003, Koyama, Y et al. 2008, Chen, Capellini et al. 2016, Shwartz, Viukov et al. 2016) also causes shortening of tibias (Capellini, Chen et al. 2017) and abnormal growth plates (Mikic, Rt et al. 2004). Yet, the cellular abnormalities related to the growth plate phenotypes were largely unknown. By implementing 3D MAPs on E16.5 *Gdf5* knockout (KO) embryos (Shwartz, Viukov et al. 2016), which already exhibited tibial deficits (Supplementary Figure 3A and 3B), we established a database of hundreds of thousands of mutant cells, which we compared to the control database.

Comparing proximal and distal growth plates of *Gdf5* KO and control tibias revealed several significantly different features. First, hypertrophic cell volume, a known determiner of growth plate height, was 24% smaller in the distal growth plate and 30% smaller in the proximal growth plate of mutant tibias relative to the control (Figure 5A). Additionally, we observed a 5% increase in the resting zone cell volume of the distal growth plate and a 10% increase in the proximal growth plate (Figure 5A). This resulted in a 2.2-fold reduction in cell growth from the resting to hypertrophic zone in the mutant distal growth plate and a 3-fold reduction in the proximal growth plate, compared to controls. In the proliferative zone, cell volumes were comparable (Figure 5A). The same trend was observed for cell surface area (Supplementary Figure 3C).

**Figure 5.**
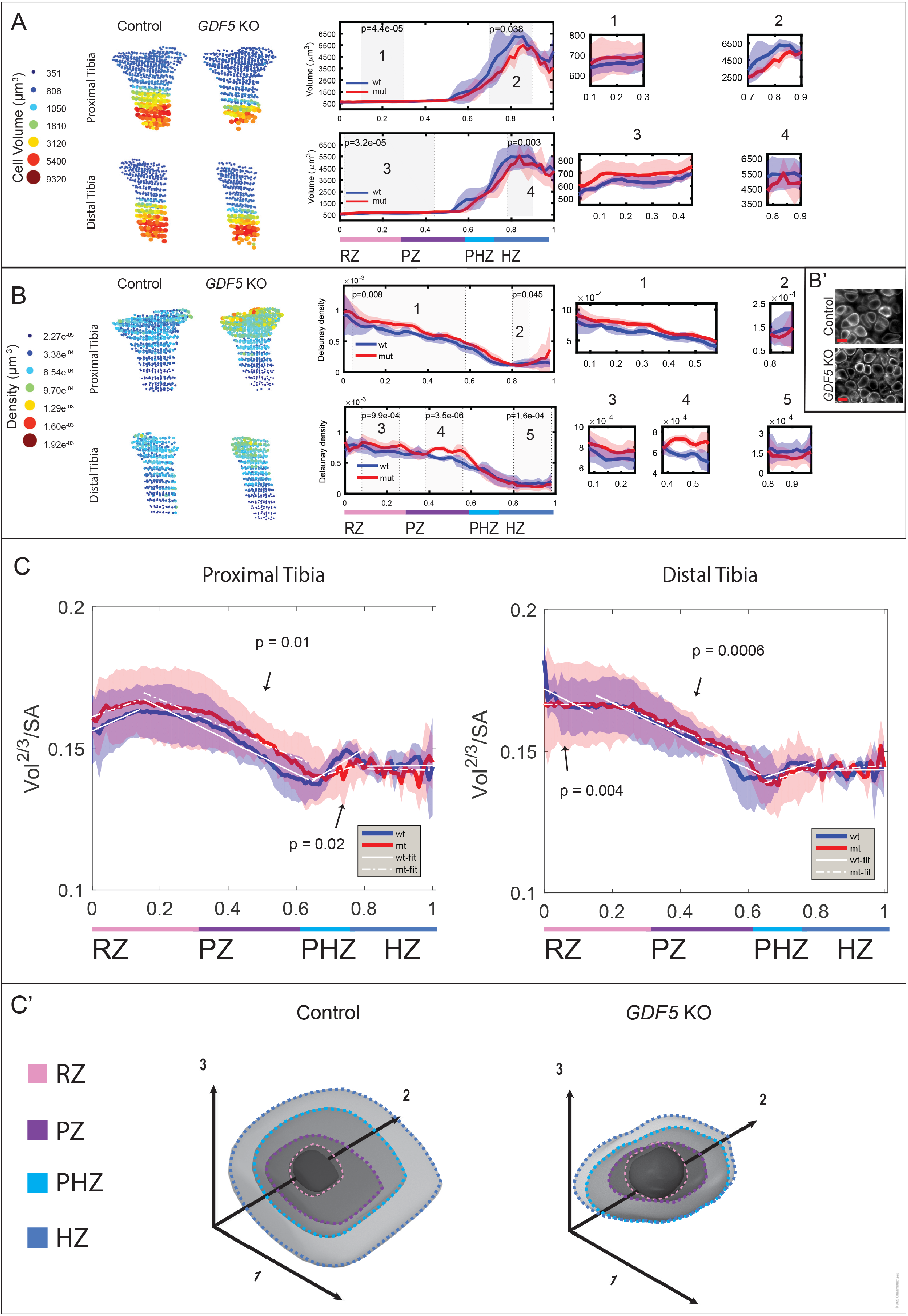
*Gdf5* regulates chondrocyte growth and morphology during differentiation. Representative 3D maps and comparative spatial profiles of control and mutant samples show that *Gdf5* KO tibias display abnormal cell volume (**A**), density (**B** and **B’**), and growth mechanisms (**C**). In the proximal (PT) and distal (DT) *Gdf5* KO growth plates (**A**), cell volume was abnormally high in the RZ (PT p = 4.4e^−05^; DT p = 3.2e^−05^) and low in the HZ (PT p = 3.8e^−02^; DT p = 3.0e^−03^). (**B**) 3D maps and spatial profiles show that the RZ and PZ of *Gdf5* KO growth plates display significantly higher cell density compared to controls (PT p = 8.0e^−03^; DT p = 9.9e^−04^ and p = 3.5e^−06^). Additionally, the HZ has lower cell density (PT p = 4.5e^−02^; DT p = 1.6e^−04^). (**B’**) Light-sheet images of cells of control and *Gdf5* KO growth plates highlights the higher density in the mutant RZ. Scale bar, 10 μm. (**C**) Spatial profile of Vol^2/3^/SA ratio with regression line fitting for each zone shows that in *Gdf5* KO growth plates, cells grew by volume dependent allometric growth in the PT RZ (slope = 0.04 p = 7.43e^−06^ R^2^ = 0.8) and isometric growth in the DT RZ (slope = 0.0 p = 0.9 R^2^ < 0.01), surface area dependent allometric growth in the PT and DT PZ (PT slope = −0.05 p = 4.75e^−29^ R^2^ = 0.93; DT slope = −0.05X p = 1.89e^−31^ R^2^ = 0.95), volume dependent allometric growth in the PT and DT PHZ (PT slope = 0.04 p = 6.44e^−04^ R^2^ = 0.58; DT slope = 0.04 p = 2.81e^−05^ R^2^ = 0.73), and isometric growth in the PT and DT HZ (PT slope = 0.0 p = 0.822 R^2^ < 0.01; DT slope = 0.01 p = 0.422 R^2^ = 0.04). The mutant growth behavior was significantly different from controls in the DT RZ (p = 4.0e^−03^), PT and DT PZ (PT p = 0.01; DT p = 6.0e^−04^), and PT PHZ (p = 0.02), where in the DT RZ the growth switched from volume dependent allometric growth to isometric growth and in the PZ and PHZ the growth mechanism stayed the same, but the growth rate was slower. (**C’**) Scheme illustrating how each cell axis changes during growth and differentiation in control and *Gdf5* KO growth plates. Abbreviations: wt = control; mut or mt = *Gdf5* KO. p-values were calculated by Student’s *t*-test between slope of 0 and slope of fitted regression line to test for isometric or allometric growth and between mean value of control (n = 5) and mutant slopes (n = 4) to find differences between controls and mutants.

Given that growth plate composition is a balance between cell number and ECM volume, and that cells were larger in the mutant resting zone, we expected to find a reduction in cell density in this region. To our surprise, cell density was higher in the resting and proliferative zones of the mutant (Figure 5B and 5B’). This suggests that mutant resting zone cells produce less ECM. Conversely, in the hypertrophic zone of the same growth plate mutant cells were less dense, had smaller cell volumes and, therefore, produce more ECM compared to controls.

Since *Gdf5* KO tibias are shorter than controls and have abnormal cell volume, surface area, and density, we next investigated if the cell growth mechanisms were impaired. Comparing the Vol^2/3^/SA between controls and mutants revealed abnormal growth behaviors in both volume-dependent and surface area-dependent allometric growth mechanisms. In the distal tibia resting zone, cells grew isometrically instead of allometrically by volume increase (Figure 5C), resulting in cell growth without a change in shape. Once they reached the proliferative zone, cells grew allometrically by surface area increase as in the wild type, but their growth rate was significantly reduced, resulting in a lesser degree of flattening (Figure 5C’). This was supported by analyzing cell sphericity, which showed that mutant cells in the resting and proliferative zones were on average 8% more spherical then control cells (Supplementary Figure 3D,3D’). This growth lag prevented mutant cells from reaching a normal size and shape even though their growth mechanisms and growth rates in the prehypertrophic and hypertrophic zones were normal (Figure 5C’).

Since the proximal tibia growth plate contributes more to bone elongation than the distal tibia (Stern, Aviram et al. 2015), and the volume defects were greater in the proximal side, we expected to see a more significant change in growth behaviors compared to the distal growth plate. As in the distal growth plate, proliferative zone cells grew at a slower rate than controls, disrupting cell flattening (Figure 5C and Supplementary Figure 3D, 3D’). Prehypertrophic zone mutant cells grew at half the rate of controls by volume-dependent allometric growth. Consequently, when cells in the hypertrophic zone grew isometrically, their final size was severely reduced and their shape impaired compared to both the mutant distal growth plate and the control growth plates (Figure 5C’). Altogether, these findings show that GDF5 plays a role in regulating both surface area- and volume-dependent growth mechanisms.

### GDF5 regulates chondrocyte polarity in the growth plate

The abnormalities we found in cell growth and flattening in *Gdf5* KO growth plates led us to examine cell organization. Indeed, we found a large variability in cell orientation along all three axes in the different zones of the mutant growth plate relative to control (Figure 6A and 6B). In view of the inability of mutant cells to flatten properly and their abnormal orientation, we suspected that there may be abnormalities in column structure. To study this possibility, we examined cross-section maps from the proliferative zone. As seen in Figure 6A(i-l) and 6E, instead of aligning their shortest axis parallel to the proximal - distal axis, as in control columns, mutant cells pointed in some areas 25-40 degrees away. In the other cell axes mutant cells varied up to 40 degrees, as compared to 15 degrees in control cells (Figure 6A(a-h)). Finally, while in controls the PC1 lines lay one on top of another, indicating that the columns are aligned along the proximal - distal axis, we observed a fan-like pattern in the mutant, supporting the possibility they form abnormal columns (Figure 6A(a-d) and 6E).

**Figure 6.**
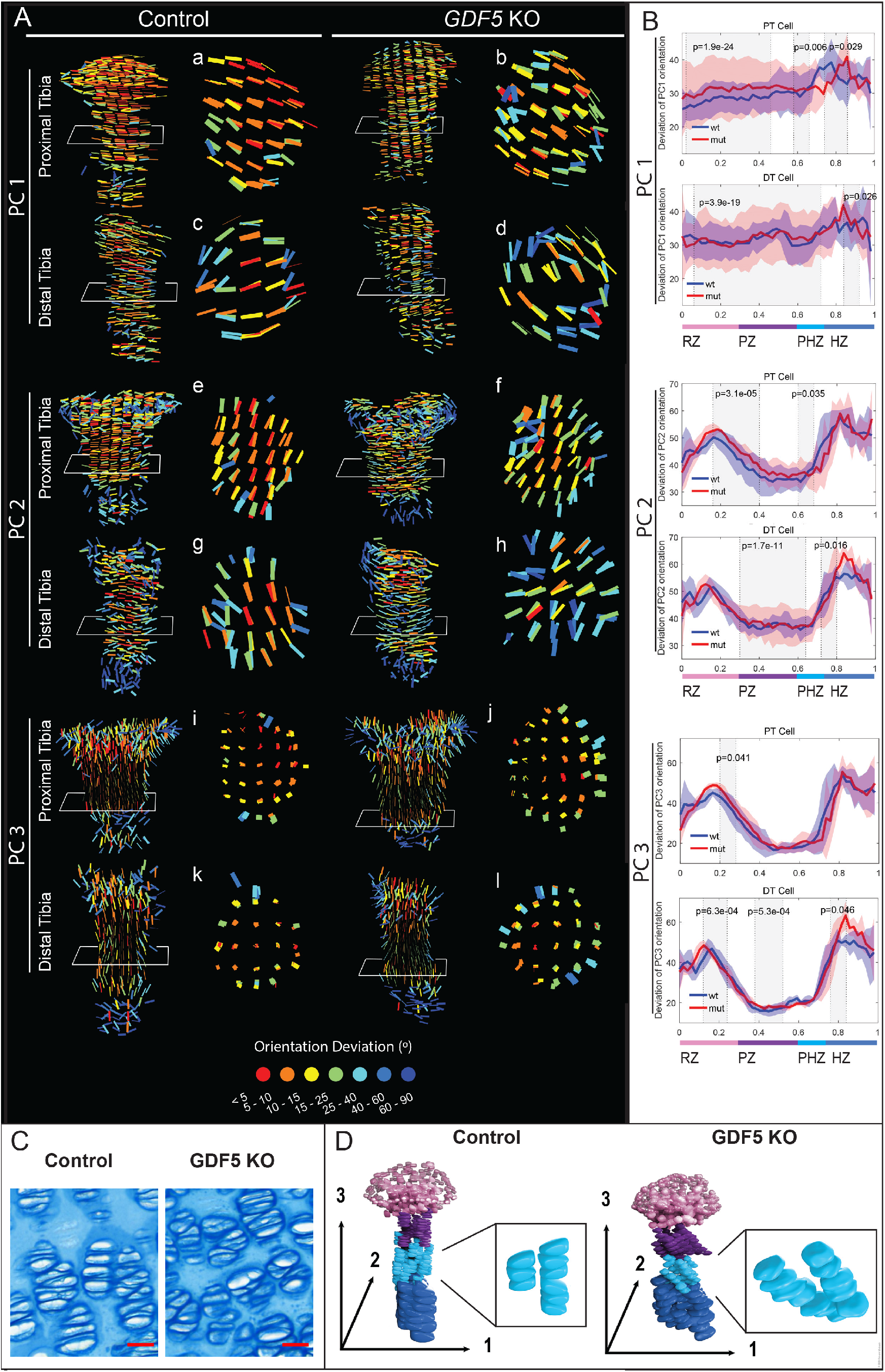
GDF5 regulates chondrocyte polarity in the growth plate. (**A**) Representative 3D maps compare cell orientation of E16.5 PT and DT growth plates between control and *Gdf5* KO mice. Cross sections through the PZ show that cells in the *Gdf5* KO growth plates orient abnormally and with high variation in PC1 and PC2 (**a-l**). The growth plates were placed such that the RZ pointed upward and the HZ pointed downward. The colormap is normalized to each sample, representing the deviation in degrees from the mean orientation of each growth plate. PC1 cross sections through *Gdf5* KO growth plates (**b** and **d**) show that lines form a fan-like pattern. This, along with the wide range of map colors, indicate that mutant cells lose their polarity along the long axis. A similar behavior was observed for PC2, and the effect was larger in the DT (**g** and **h**). Cells in control growth plates aligned their shortest axis (PC3) along the proximal-distal axis of the bone within a range of 5 degrees (**i** and **k**). Strikingly, cells in *Gdf5* KO growth plates were polarized in this axis, but they deviated by 25-40 degrees (**j** and **l**). (**B**) Spatial profiles show how PC 1, 2, and 3 differ between control and *Gdf5* KO growth plates. PC1 cell orientation was significantly more variable in the mutant in multiple locations throughout all zones in both PT and DT growth plates (PT, p = 1.9e^−24^ and 6.0e^−03^ and 2.9e^−02^; DT, p =3.9e^−19^ and 2.6e^−02^). PC2 cell orientation was significantly more variable only in PHZ of the PT (p = 3.5e^−02^) and in the PZ and HZ of the DT (p = 1.7e^−11^ and 1.6e^−02^). PC3 cell orientation displayed the least variability compared to the other axes. In the mutant PT, the variability was significantly lower in the RZ (p = 4.1e^− 02^). The DT growth plate displayed more significant changes. *Gdf5* KO growth plates were less variable in the RZ and PZ (p = 6.3e^−04^ and p = 5.3e^−04^), but more variable in the HZ (p = 4.6e^−02^). (**D**) Alcian blue staining of PZ from control and *Gdf5* KO tibias shows abnormal column structure in the mutant. Scale bar, 20 μm. (**E**) 3D model of cell orientation in control and *Gdf5* KO growth plates. Insets highlight columns in the proliferative zone. Abbreviations: wt, control; mut, *Gdf5* KO. p-values were calculated by Student’s *t*-test between standard deviations (shaded regions) of control (n=5) and mutant samples (n=4).

To validate these statistical results, we performed histological analysis on control and mutant growth plates at P6, when the column structure is already well established (Figure 6C). Alcian blue staining clearly showed that mutant columns were disorganized compared to controls. Overall, we observed defects in cell orientation and column formation that may impair growth plate function, leading to the observed growth defects of *Gdf5* KO tibias.

## Discussion

The tight connection between cell growth and morphogenesis on one hand, and organ development and function on the other, highlights the need to study the mechanisms that underlie this association. For that, it is necessary to characterize in detail the 3D morphology of cells in the spatial context of the tissue. Yet, high cell number, organ size and dense extracellular matrix have thus far limited the ability to conduct such analyses in mammalians.

In this work, we established a new pipeline called 3D MAPs, which enabled us to overcome these limitations and create complex multiresolution datasets of the 3D morphology of growth plate cells throughout their differentiation. By analyzing these datasets, we show that chondrocytes use both allometric and isometric growth strategies during differentiation in a cell axis-specific manner. Additionally, 3D MAPS revealed a striking conservation in the longest cell axis orientation across all zones of the growth plate. Finally, we analyzed tibias of mice lacking *Gdf5* and identified multiple abnormalities in cell growth, shape and organization, which can explain the shortening of these bones and uncover a new regulatory role for GDF5 in growth plate biology.

Nearly a century ago, Stump proposed that “The growth impulse affecting the shape, size, and form of the bone is inherent in the cartilage cells, both in diaphysis and epiphysis.” (Stump 1925). Indeed, the morphology of growth plate chondrocytes during endochondral ossification and, in particular, their flattening during column formation and their swelling during terminal differentiation have attracted much attention over the years (Stump 1925, Dodds 1930, Breur, Vanenkevort et al. 1991, Wilsman, Farnum et al. 1996, Breur, Md. et al. 1997, Wilsman, Bernardini et al. 2008, Li and Dudley 2009, Cooper, Oh et al. 2013, Romereim, Conoan et al. 2014, Li, Li et al. 2017, Cooper 2019). Although these studies have greatly promoted our understanding of this process, the full sequence of morphological changes that differentiating chondrocytes undergo was largely missing. Since the growth plates regulate bone morphology and elongation, revealing this sequence is imperative for appreciating the complexity of the process and uncovering the mechanisms that underlie it.

To achieve that, it is necessary to obtain 3D information on cell shape throughout the differentiation process. Although there have been several attempts to study 3D morphology of cells in the growth plate (Cruz-Orive and Hunziker 1986, Guilak 1995, Guilak, Ratcliffe et al. 1995, Amini, Veilleux et al. 2010, Amini, D. et al. 2011), these studies focused on comparing small discrete regions from each zone, which did not produce the statistical power needed to understand the exact contribution of each morphological change to growth plate function.

3D MAPs offers solutions to the multiple problems and challenges in studying 3D cell morphology in the context of the entire tissue. First, it provides a modular tool kit with which to image, register, segment, and quantify morphological properties of hundreds of thousands of cells in a time efficient manner. Multiresolution capabilities allow the user to shift from single-cell to tissue level to identify morphological patterns amongst thousands of cells within the tissue. Moreover, 3D MAPs allows quantification of 17 different morphological parameters, thereby enabling one to measure the particular contribution of each differentiation step to cell growth and morphogenesis. Finally, while studying mutants in which growth plate activity is affected, it is possible to identify the exact cellular abnormality and the stage at which chondrocyte differentiation deviates from the normal process. This feature is particularly important as it can point to the molecular mechanism underlying the phenotype.

Cells can grow either isometrically, where only cell size changes, or allometrically, where both size and shape change. 3D MAPs analysis revealed that differentiating chondrocytes employ both strategies. During resting, proliferating, and prehypertrophic stages cells grow allometrically, whereas the final growth stage in the hypertrophic zone is isometric. Interestingly, we found that allometric growth can be driven by increase of either surface area or volume. Another interesting finding is that allometric growth can be axis-specific. For example, in the resting zone allometric growth was achieved by swelling along the shortest and second longest cell axes. Conversely, in the proliferative zone it involved expansion of the cells along the longest and second longest axes and shortening of the shortest axis. Finally, in the prehypertrophic zone, cells kept the longest axis constant while swelling preferentially along the remaining two axes. Altogether, our results show that chondrocyte growth strategies are dynamic and complex. Moreover, they predict the existence of multiple mechanisms that are needed to regulate this complex process. It is interesting to note that although resting zone and hypertrophic cells are 9-fold different in volume, they share the same aspect ratio. This suggests that in principle, resting zone cells could have reached the same volume and shape as hypertrophic cells through isometric growth. Thus, the need for multiple strategies of allometric cell growth is either fundamental to growth plate biology, or a secondary consequence of the innate structure of the growth plate.

In many tissues such as the epithelium, pancreas, and liver, cell size was shown to regulate cell proliferation in order to maintain population size (Ginzberg, Kafri et al. 2015). This is achieved by only allowing cells of a certain size to enter S phase and divide (Dolznig, Grebien et al. 2004). Interestingly, growth plate cells increased their size up to 5-fold while being proliferatively active, moving from the resting to the prehypertrophic zone. This suggests that chondrocytes in each zone reset their cell size measurement mechanism in order to allow proliferating chondrocytes to increase their size.

Cell morphology and tissue architecture are tightly connected (Irvine and Wieschaus 1994, Hogan 1999, Lecuit and Lenne 2007, Martin, Kaschube et al. 2009). For decades, researchers have been fascinated with growth plate column formation, as it represents a transition from disorder to order (Dodds 1930, Li and Dudley 2009, Romereim, Conoan et al. 2014, Li, Li et al. 2017). Because this process was studied in 2D, orientation of the shortest cell axis and its relation to column formation received virtually all the attention. Using 3D MAPS, we found that the axis that orients along the M-L axis, which previous studies suggested as the cell’s major axis (Li and Dudley 2009, Li, Li et al. 2017), is only the second longest axis of the chondrocytes. The longest cell axis, which is more than twice the length of the second longest, is oriented towards the D-V axis. Surprisingly, we discovered that unlike the two other axes, this orientation was conserved in all growth plate zones. Over the years, several mechanisms were shown to influence chondrocyte morphology and orientation. Polarity-inducing molecules, such as members of PCP pathway and integrin molecules are involved in chondrocyte shape and orientation (Aszodi, Hunziker et al. 2003, Li and Dudley 2009, Gao, Song et al. 2011, Li, Li et al. 2017). Additionally, ECM stiffness (Prein, Warmbold et al. 2016) and mechanical signaling contribute to chondrocyte polarity (Shwartz, Farkas et al. 2012). Although the reason for chondrocytes to align their longest axis along the D-V bone axis is unknown, we speculate that it may provide a structural advantage by increasing the stability of the growth plate during development.

GDF5 plays important roles in joint development and disease (H. and Lee 1973, Storm, T.V. et al. 1994, Mikic, Rt et al. 2004, Koyama, Y et al. 2008, Shwartz, Viukov et al. 2016, Capellini, Chen et al. 2017). Notably, it was shown to be expressed in the outer layer of the growth plate (Capellini, Chen et al. 2017) and to have a role in bone growth, as humans with Grebe syndrome, a recessive mutation in *Gdf5*, as well as mice lacking *Gdf5* displayed shortening of limbs and misshapen bones (Grebe 1952, Storm, T.V. et al. 1994, Mikic, Rt et al. 2004, Basit, Naqvi et al. 2008, Chen, Capellini et al. 2016, Capellini, Chen et al. 2017). However, the cellular components underlying the shortening and misshaping of *Gdf5* mutant bones have not been described. We identified in growth plates of *Gdf5*-null mice several abnormalities in cell growth, morphology and orientation that resulted in aberrant allometric and isometric growth. Chondrocytes in the resting zone were abnormally larger in the mutant; yet, surprisingly, despite this size advantage, in the prehypertrophic zone they failed to swell fast enough, resulting in significantly smaller hypertrophic cells. This growth defect was amplified by a lower cell density in the hypertrophic zone. Additionally, a reduced surface area-dependent allometric growth rate in the proliferative zone, combined with more spherical resting zone cells, impaired cell flattening, which most likely contributed to misorientation of cells along all three cell axes. For example, cells in the proliferative zone deviated 15 degrees more compared to controls in all three axes, likely causing aberrant column formation in this zone. Given the known connection between hypertrophic zone cell size and bone length (Breur, Vanenkevort et al. 1991, Breur, Md. et al. 1997, Cooper, Oh et al. 2013), as well as the link between cell polarity in the proliferative zone and misshapen bones (Li and Dudley 2009, Gao, Song et al. 2011, Romereim, Conoan et al. 2014, Li, Li et al. 2017), these cellular phenotypes can together explain the dysplastic *Gdf5* KO tibias. These findings raise many questions, such as whether the effect of GDF5 is autonomous or nonautonomous, as well as the identity of the molecular mechanism that mediates the role of GDF5 in regulating chondrocyte growth, shape and organization.

In summary, this work introduces a pipeline that allows in-depth profiling of cell growth, morphology, and organization in the spatial context of a mammalian tissue. 3D MAPs provides a means to explore how the growth plate works and to identify the mechanisms that regulate it. As a proof concept, analysis of mutant mice identified GDF5 as a regulator of growth plate activity. This new pipeline has the potential to change the way we study growth plate biology.

## Methods

### Animals

For genetic labeling of chondrocyte lineage in control and *Gdf5* KO mice, *mTmG:Col2a1-Cre:Gdf5-CreER* control and mutant mice were used in all analyses. To generate *mTmG:Col2a1-Cre:Gdf5-CreER* mice, animals homozygous for *mTmg* (Jackson Laboratories) were crossed with *Col2a1-Cre* mice (Jackson Laboratories). Then, *mTmG:Col2a1-Cre* mice were crossed with mice heterozygous for *Gdf5-CreER* (Shwartz, Viukov et al. 2016). Embryos were dissected in cold phosphate buffered saline (PBS), fixed overnight at 4°C in 1% paraformaldehyde (PFA), washed in PBS and stored at 4°C in 0.5 M EDTA (pH 8.0, Avantor Performance Materials) with 0.01% sodium azide (Sigma). In all timed pregnancies, the plug date was defined as E0.5. For harvesting of embryos, timed-pregnant female mice were sacrificed by CO_2_ exposure. Embryos were sacrificed by decapitation with surgical scissors. Tails were visualized with fluorescent binoculars for genotyping when possible; alternatively, tail genomic DNA was used for genotyping by PCR. All animal experiments were pre-approved by the Institutional Animal Care and Use Committee (IACUC) of the Weizmann Institute.

### Histology

E18.5 or P6 tibias were fixed in 4% PFA overnight, dehydrated to 100% ethanol (dehydration sequence: 25%, 50%, 70%, and twice in 100% for one hour each), and embedded in paraffin. 7-μm-thick paraffin sections were stained with H&E or Alcian Blue and collected onto slides. Slides were mounted with entellan and digitized with a Pannoramic scanner (3DHISTECH) and images were exported with QuPath software (Bankhead, Loughrey et al. 2017).

### Tissue clearing

E16.5 tibias and ulnas from *mTmG:Col2a1-Cre:Gdf5*-*CreER* control and mutant mice were cleared using the PACT-deCAL technique (Yang, Treweek et al. 2014, Treweek, Chan et al. 2015). Shortly, decalcified samples were washed in PBS, then embedded into a hydrogel of 4% (wt/vol) acrylamide in 1x PBS with 0.25% thermal initiator 2,2’-azobis[2-(2-imidazolin-2-yl)propane]dihydrochloride (Wako, cat. No. VA-044). The hydrogel was allowed to polymerize at 37°C for 3 hours. The samples were removed from the hydrogel, washed in PBS, and moved to 10% SDS with 0.01% sodium azide, shaking at 37°C for 4 days, changing the SDS solution each day. Samples were washed four times with 1x PBST (PBS + 0.1% Triton X-100 + 0.01% sodium azide) at room temperature (RT) over the course of a day. To label nuclei, samples were submerged in 8 μg/ml DAPI in 1x PBST gently shaking overnight at RT. Samples were washed again with four changes of 1x PBST, and the refractive index (RI) of the sample was brought to 1.45 by submersion in a refractive index matching solution (RIMS) consisting of Histodenz (Sigma) and phosphate buffer, shaking gently at RT for 2-3 days. Samples were embedded in 1% low gelling agarose (Sigma) in PBS, in a glass capillary (Brand, Germany). Embedded samples were submerged in RIMS and protected from light at RT until imaging.

### Soft tissue micro-CT

P12 control and *Gdf5* KO tibias were dissected, fixated overnight in 4% PFA in PBS, and dehydrated to 100% ethanol (dehydration sequence: 25%, 50%, 70%, and twice in 100% for one hour each). According to (Metscher 2009), samples were soaked in 2% iodine in 100% ethanol solution (Sigma) for 48 hours at 4°C. Tissue was washed in 100% ethanol twice for 30 minutes each prior to scanning. Samples were then mounted in 100% ethanol and scanned by MicroXCT-400 (Xradia) at 30 kV and 4.5 W with the Macro-70 lens.

### Light-sheet microscopy

Cleared samples were imaged using a light-sheet Z1 microscope (Zeiss Ltd) equipped with 2 sCMOS cameras PCO- Edge, 10X illumination objectives (LSFM clearing 10X/0.2) and Clr Plan-Neofluar 20X/1.0 Corr nd=1.45 detection objective, which was dedicated for cleared samples in water-based solution of final RI of 1.45. A low-resolution image of the entire tibia was taken with the 20x Clarity lens at a zoom of 0.36. To acquire higher resolution images of the proximal and distal growth plates, multiview imaging was done with the same lens at a zoom of 2.5 resulting in x,y,z voxel sizes of 0.091 μm, 0.091 μm, 0.387 μm. The DAPI and Col2Cre-mGFP channels were acquired with Ex’ 405 nm Em’BP 420-470 and Ex’ 488 nm Em’ BP 505-545 lasers at 2.4% and 3% laser powers, respectively. Light-sheet fusion of images was done in Zen software (Zeiss). Stitching of low-resolution images was done in ImarisStitcher.

### Growth plate segmentation

Prior to segmentation, images were down-sampled in the X,Y direction using the Downsample plugin in Fiji (Schindelin, Arganda-Carreras et al. 2012) resulting in x,y,z voxel size of 0.194 μm, 0.194 μm, 0.387 μm and saved as TIFF files, which were then converted to VFF file format in MatLab (S1 data). Non-cartilaginous regions were masked using Microview 2.1.2 (GE Healthcare). Shortly, spline contours were drawn around the growth plate throughout each z-stack, and the pixels outside the resulting 3D region of interest were reassigned an intensity value of 0 (S2 data).

### Nucleus segmentation

To automatically segment fluorescently-labeled nuclei in the 3D images, we performed a two-step procedure. In the first step, seed points that were roughly located in the center of the nuclei were detected using a Laplacian-of-Gaussian-based (LoG) approach as described in (Stegmaier, Otte et al. 2014). In brief, the 3D input images were filtered with a LoG filter using a standard deviation that was empirically tuned to the radius of the objects of interest. We used standard deviations of σ = 12 for RZ, PZ and PHZ nuclei and of σ = 35 for HZ nuclei. Subsequently, local intensity maxima were extracted from the LoG-filtered image and reported as potential nuclei centers. For each potential seed point, we compute the mean intensity in a 4×4×4 voxel-wide cube surrounding the centroid. In order to minimize the number of false-positive detections, only seed points with a mean intensity larger than the global mean intensity of the entire LoG-filtered image were kept for further processing. In a final step, we used a seeded watershed algorithm to perform the segmentation of the nuclei in a Gaussian-smoothed 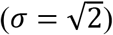 version of the intensity-inverted raw input image. The detected seed points were used to initialize the seeded watershed algorithm and we artificially added a background seed located at the border of each image snippet to separate the centered nucleus from the surrounding background. The segmentation was performed separately for each nucleus and in parallel, *i.e.*, small 3D image patches surrounding the seed points were processed concurrently using multiple cores of the CPU. Segmentation results of the individually processed patches were then combined to form a full-resolution segmentation image containing the final result with a unique integer label for each of the nuclei that was used for further quantification and morphological analyses. All image analysis pipelines were implemented using the open-source software tool XPIWIT (Bartschat, Hübner et al. 2015) and executed on a Windows Server 2012 R2 64-bit workstation with 2 Intel(R) Xeon(R)CPU E5-2690 v3 processors, 256 GB RAM, 24 cores and 48 logical processors.

### Cell segmentation

To segment fluorescently labeled cell membranes, we performed a manual contrast adjustment of the input images and extracted cell segments using a morphological watershed algorithm (Beare and L. 2006). Due to intensity variation in different regions of the image, we processed individual tiles separately and manually tuned the watershed starting level to obtain a good segmentation, i.e., all local minima below the specified level were used to initialize the catchment basins for the watershed regions. All image analysis pipelines were implemented using the open-source software tool XPIWIT (Bartschat *et al.*, 2016) and executed on a Windows Server 2012 R2 64-bit workstation with 2 Intel(R) Xeon(R)CPU E5-2690 v3 processors, 256 GB RAM, 24 cores and 48 logical processors.

### Nucleus and cell feature extraction

Following image segmentation, images were resaved with multiple thresholds in order to avoid merging of neighboring objects during image binarization (S3 data). Then, cell and nucleus volumes were measured using 3D manager (Ollion, Cochennec et al. 2013) for each growth plate zone of each of the five growth plates, in order to set a minimal and maximal volume range to further clean the data (Supplementary Table 2). Finally, cells and nuclei overlapping the image borders were removed. To extract morphological features, each nucleus and cell were first converted from a binary volume into a triangulated mesh by applying Gaussian smoothing filter (sd = 0.5 pixels) and extracting the iso-surface at the iso-value 0.1. From the triangulated mesh of each object (nucleus or cell), the following features were extracted: surface area, as the sum of the areas of all mesh faces; volume, using the convergence theorem (a.k.a. Gauss’s theorem or Ostrogradsky’s theorem (Gauß 1813)); sphericity, calculated as:

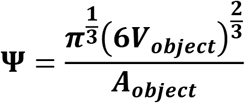

where ***V***_***object***_ is the volume of the object and ***A***_***object***_ is the surface area (Wadell 1932); object orientation of each principal axis, defined as the direction of the first (PC 1), second (PC 2), or third (PC 3) principal components of the masked region of the object in real distance units. PC1 is the largest, PC2 the second largest, and PC3 is the smallest principal components. Feature extraction was done on an Intel® Xeon® CPU E5-1620 v4 @ 3.50GHz, 32.0 GB RAM, Windows 10 64 bit, with MatLab version 2017b (The MathWorks, Inc., Natick, Massachusetts, USA).

### Segmentation validation

To validate the segmentation measurements, an E16.5 tibia from a sample littermate was dissected and cleared using PACT-deCAL, stained with DAPI, mounted in RIMS on a glass slide with a shallow well, and covered with a glass coverslip. 75-μm z-stacks from areas in the RZ, PZ, PHZ, and HZ were acquired with x,y,z voxel sizes of 0.1 μm, 0.1 μm, 0.5 μm. Confocal images were manually segmented (cells and nuclei: n_RZ_ = 20; n_PZ_ = 20, n_PHZ/HZ_ = 40) in Microview software and volume and surface area were calculated for each. Then, volume and surface area of automatically segmented cells and nuclei (cells: n_RZ_ = 17136; n_PZ_ = 6698, n_PHZ/HZ_ = 4174; nuclei: n_RZ_ = 56812; n_PZ_ = 12435, n_PHZ/HZ_ = 12153) from comparable regions in the growth plate were compared to the manual segmentations by a chi-square distance between histograms test (Porter 2008). For this, we performed independent sampling with a pool size of 20 for each zone, and repeated the procedure 10,000 times to ensure the sampling covers the variability in these zones. The average bins of the 10,000 iterations were plotted as a histogram and a two-sample chi-square distance test was conducted between non-zero bins of the automatic and manual segmentations.

### Segmentation error calculation

To calculate segmentation error from all samples, we calculated the mean ratio of correctly segmented objects (nuclei or cells) out of the total object number in every image across all samples (n = 20 growth plates). For this, we counted the number of objects in a 2D optical section every 23 μm along the Z axis, to avoid counting the same object twice (S4 data). We then divided the number of automatically segmented objects in matching 2D sections to the number counted from the raw images to get the segmentation error for each image (S5 data). We considered a nucleus or cell to belong to a given 2D slice if its centroid was within 10 z planes (3.87 μm) from the given slice. We then averaged the errors within and across all samples (Supplementary Figure 1C) to calculate the mean error for nuclei and cells.

### Growth plate registration

All growth plates (PT, DT, and DU) were registered based on the outer surface of the RZ. For this, the mean of nuclear centroids were fixed to the origin of each growth plate sample. To remove bias due to heterogeneous density of centroids, only centroids located on the surface boundary points were used. Then, a bounding volume of each growth plate was extracted using the MatLab function alphaShape polygons. MatLab functions AlphaTriangulation and freeBoundary were used to extract the vertices from the outer surface of the growth plates and eigenvectors were calculated from the covariance of the vertices. This eigenvector matrix was used for the growth plate registration. We checked manually that distal, proximal, medial, lateral, dorsal and ventral parts of the RZ matched between growth plates. Manual multiplication of 180-degree rotation matrix was used to correct the eigenvectors if needed. The coordinates of the registered growth plates were calculated by multiplying the eigenvectors with the centroids.

### 3D morphology maps of growth plates

To visualize the spatial distribution of cellular or nuclear morphology within each growth plate, data were first stitched back together in MatLab using the stage coordinates and scaling of each image. To place each voxel in a global coordinate system, we added the coordinates of each stack to every voxel in the stack, thereby reconstructing the entire bone. Growth plates were then registered based on manual alignment of the condyles in the RZ end of the growth plate. To highlight large patterns while averaging out small differences between individual nuclei, we represented nuclear and cellular features at a coarse-grained level. This representation was computed by first defining a regular spatial grid over the data. Each element of the grid was defined as a 75 × 75 × 75 μm cube containing up to 532 nuclei, and 255 cells and at least 178 nuclei and 119 cells. Within each cube, we averaged the nucleus and cell volume, surface area and sphericity and used the Delaunay tessellation field estimator at a resolution of 2 μm^3^ to compute the nuclear density for each cube (Schaap 2007). Since most growth plate cells at E16.5 have only one nucleus, nuclear density was used to infer cell density as well. In addition, for each cube, we computed the occupation, defined as the sum of nuclear or cellular volumes divided by the volume of the cube, N/C ratio and all other features as shown in Supplementary Table 1. Then, using MatLab’s jet color map, we represented the characteristics of each cube on the grid by drawing spheres whose radii and colors are proportional to the computed values and whose centers correspond to the average position of the nuclei or cells within the bin.

### Orientation maps

The orientation of the three axes of a cell or a nucleus is described by the first, second, and third principal components of a PCA performed on the point cloud corresponding to the segmented shape of the cell or nucleus. When computing the principal component, the sign of the vector was attributed arbitrarily. Before averaging the orientations in a cube, we first constrained the orientations to be in the same hemisphere to avoid artificial bias. This was performed by considering all the orientations and their opposites, leading to 2N vectors, where N is the number of objects in the cube. We then performed a PCA on the set of orientation vectors to determine the main direction. The N orientation vectors corresponding to the hemisphere representing the main direction were extracted by considering only the ones having a positive dot product with the main direction. The main orientation was computed over the N selected orientation vectors as a standard average over the direction cosines (Mardia 2014). The spherical variance was computed as the dispersion around the mean direction (Mardia 2014). For each cube, a line was drawn oriented and color-coded according to the deviation from the mean orientation of the whole growth plate, and whose thickness was proportional to the spherical variance. Finally, to summarize the distribution of orientations in a 3D map, we measured the difference of angle variation between mean orientation vectors obtained for each of the bins with mean orientation vectors obtained for the whole growth plate. A seven binned rainbow colormap (red to blue) was assigned to show small to large variations in angles.

### Spatial profiles of morphology features

To visualize quantitatively the spatial profile of each of the computed features and identify gradients along the growth plate, the entire growth plate was divided into 50 equally spaced bins along the P-D axis. For each bin, the mean value of individual cell or nucleus features were computed from the cells/nuclei whose centroid fell in this bin. Finally, bins were plotted with the X-axis representing the P-D axis and Y-axis representing the mean value of the feature among all samples for each bin.

### *t*-test of the means and standard deviations of morphological features

To detect significantly different regions along the P-D axis of two groups of growth plates, a *t*-test on the spatial profile of a given feature was performed. The regions were tested by two ways, first using the mean value of features and second using the standard deviation of features between groups; i.e. five control and five mutant samples. P-value ≤ 0.05 indicates statistical significance. The *t*-tests were performed as follows: We chose the window length of 5 and checked for the significant regions in the window of bins [i to i+5], where i index ranges from 1 to 45. All the windows the bins of which had p ≤ 0.05 were merged and the final p-value was recalculated on these merged bins, showing the significant regions in the graph.

### Linear regression of Vol^2/3^/SA graphs

Four linear regressions were fit to data in RZ, PZ, PHZ, and HZ using the MatLab function POLYFIT. The boundary between zones in the x-range was adjusted manually, such that each growth plate used the same range of data. The slope and intercept value of the linear regression are mentioned in the figure or legend (Figures 3 and 5) and the fitted line was plotted using POLYVAL function. If actual and fitted data is y and yfit, respectively, then coefficient of determination R^2^ is

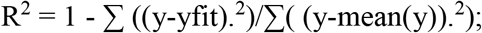

and standard error of slope coefficient is

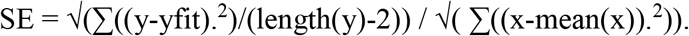

p-value of the slope was calculated using Student’s *t*-test cumulative distribution function

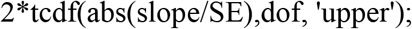

where the slope is the slope coefficient and dof is the degrees of freedom in the given data.

## Acknowledgements

We thank Nitzan Konstantin for editorial assistance and members of the Zelzer lab for their advice and encouragement throughout this project. We thank the de Picciotto Cancer Cell Observatory in memory of Wolfgang and Ruth Lesser, Weizmann Institute of Science, for providing LSFM infrastructure, Rada Massarwa and Jacob Hanna from the Department of Molecular Genetics, Weizmann Institute, for providing *mTmg* mice, Ofra Golani from the MICC Cell Observatory and Kiril Kogan from the Bioinformatics Unit, Weizmann Institute, for support and discussions regarding segmentation analysis, Ron Rotkopf from the Bioinformatics Unit for consulting us on statistics, Tali Wiesel from the Graphic Design Department at the Weizmann Institute of Science for her help with graphics, and Chinami Michaels for her help creating schemes and an animation of our findings.

## Author Contributions

S.R. designed and carried out experiments and analyses and wrote the manuscript. T.S., P.V., and A.A. designed and carried out analyses. J.St. designed analyses. J.Sv. assisted with data preprocessing. Y.A. assisted with benchmark imaging. E.Z. designed and supervised experiments and analyses and wrote the manuscript. All authors reviewed the manuscript.

## Competing interests

The authors declare no competing interests.

## Funding

This study was supported by grants from the National Institutes of Health (NIH, #R01 AR055580), European Research Council (ERC, #310098), the Israel Science Foundation (ISF, #345/16), the David and Fela Shapell Family Center for Genetic Disorders, and the Estate of Bernard Bishin for the WIS-Clalit Program (to E.Z), the WIN program between Princeton University and the Weizmann Institute (to P.V.), and from the German Research Foundation (DFG-MI1315/4-1; to J.S).

## Supplementary Information

**Supplementary Table 1.**
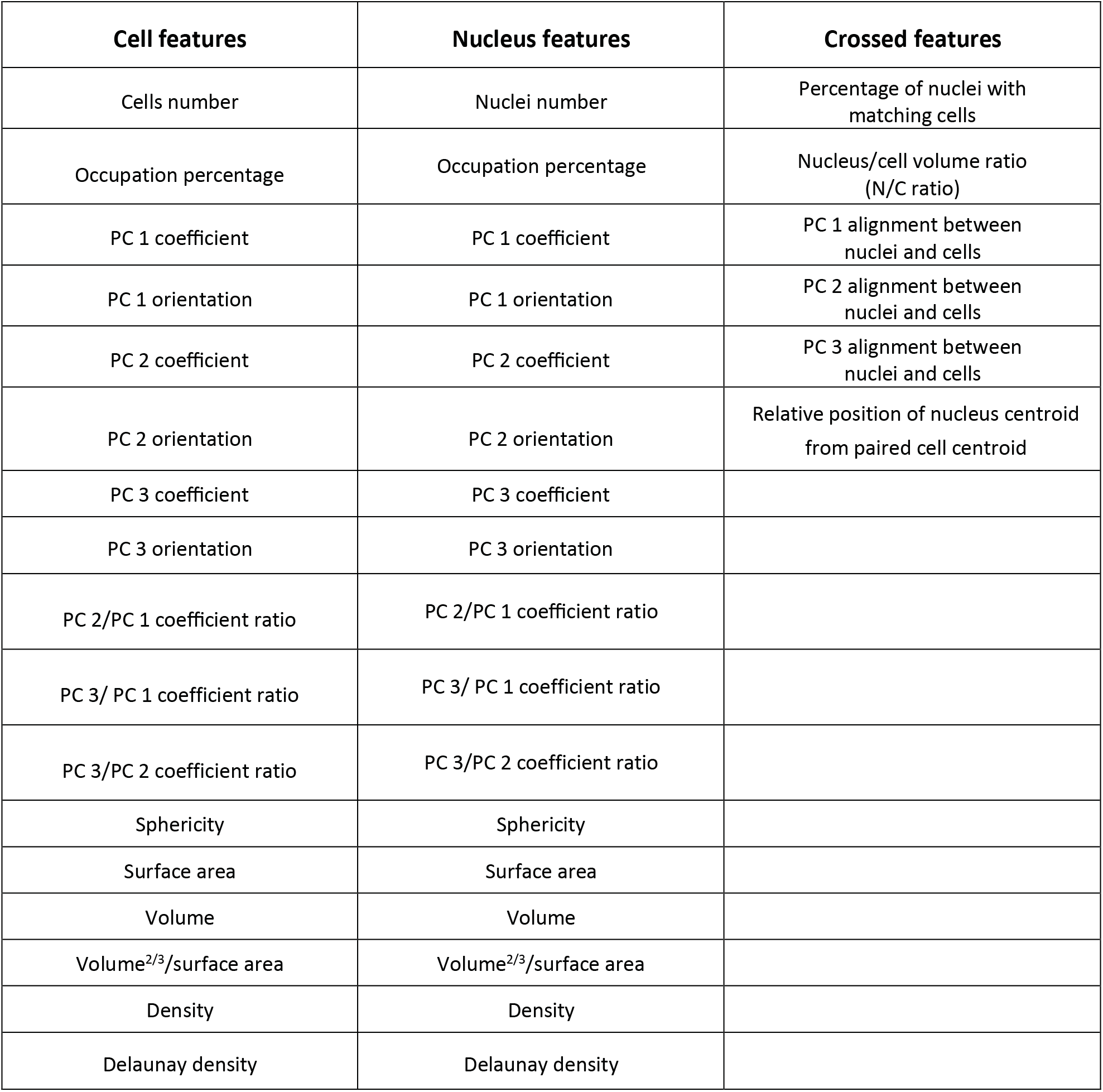
3D MAPs extracted features.

**Supplementary Table 2.**
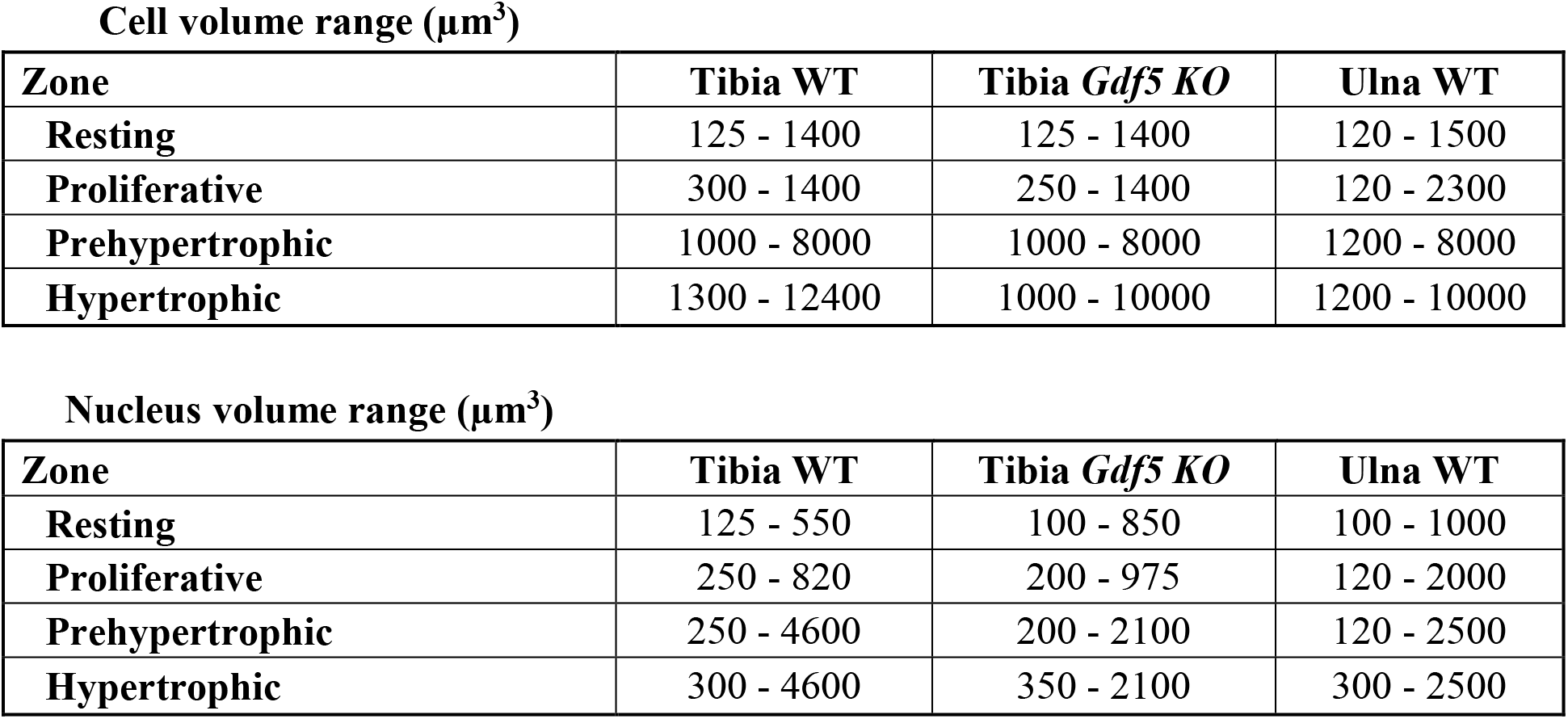
Cell and nucleus volume ranges. To eliminate erroneous cell and nucleus segmentations, we excluded from the analysis objects that had a volume outside of a given range. These ranges were set by manually selecting correctly segmented cells and nuclei from each zone and calculating their volumes in 3D morphology manager (Ollion, Cochennec et al. 2013).

**Cell volume range ( $\mu\text{m}^3$ )**
| **Zone** | **Tibia WT** | **Tibia *Gdf5* KO** | **Ulna WT** |
| --- | --- | --- | --- |
| **Resting** | 125 - 1400 | 125 - 1400 | 120 - 1500 |
| **Proliferative** | 300 - 1400 | 250 - 1400 | 120 - 2300 |
| **Prehypertrophic** | 1000 - 8000 | 1000 - 8000 | 1200 - 8000 |
| **Hypertrophic** | 1300 - 12400 | 1000 - 10000 | 1200 - 10000 |

**Supplementary Table 2. Cell and nucleus volume ranges.**
| **Zone** | **Tibia WT** | **Tibia *Gdf5* KO** | **Ulna WT** |
| --- | --- | --- | --- |
| **Resting** | 125 - 550 | 100 - 850 | 100 - 1000 |
| **Proliferative** | 250 - 820 | 200 - 975 | 120 - 2000 |
| **Prehypertrophic** | 250 - 4600 | 200 - 2100 | 120 - 2500 |
| **Hypertrophic** | 300 - 4600 | 350 - 2100 | 300 - 2500 |

**Supplementary Figure 1.**
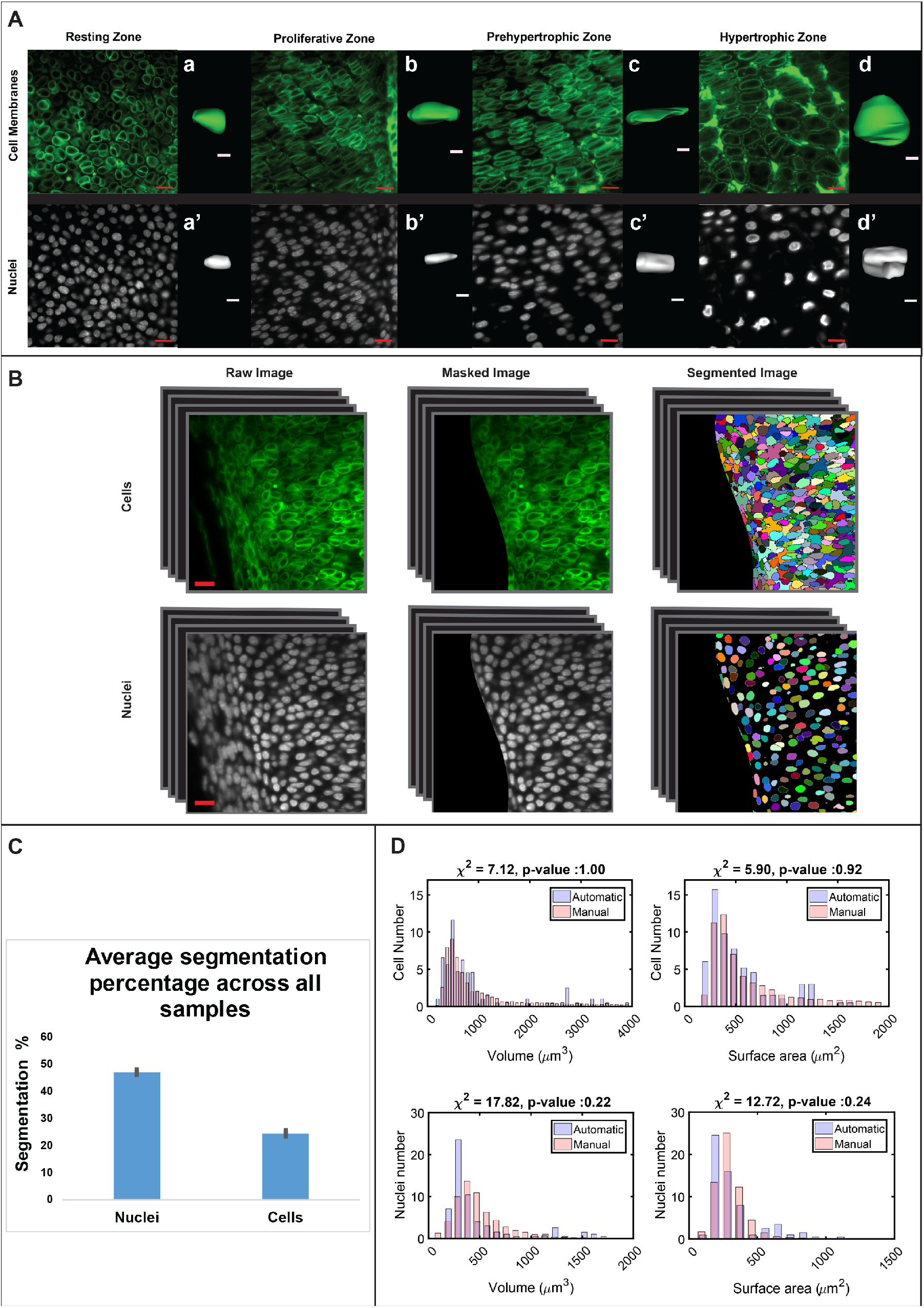
Data acquisition and segmentation. (**A**) Light-sheet images of cells and nuclei from the resting (RZ), proliferative (PZ), prehypertrophic (PHZ), and hypertrophic zones (HZ) were captured at 2.5X optical zoom with a 20X clarity lens of a Zeiss Lightsheet Z.1 microscope. Cell membranes were endogenously labeled by using *mTmG:Col2a1-Cre* mice and nuclei were stained with DAPI. Scale bars: 20 μm. (a-d) 3D surface rendering of cells from the RZ, PZ, PHZ, and HZ. Scale bars: 5 μm. (a’-d’) 3D surface rendering of nuclei from the RZ, PZ, PHZ, and HZ. Scale bars: 5 μm. (**B**) Cells and nuclei undergo semi-automatic segmentation in Microview and XPIWIT. For both, the 3D raw image is masked, where the cells and nuclei not belonging to the growth plate are removed. Then, the masked image undergoes automatic segmentation for cells and nuclei with specially designed algorithms in XPIWIT to produce a 3D segmented image. Each color in the segmented image represents an individual object. Scale bars: 20 μm. (**C**) The average segmentation error was calculated for all growth plate samples by calculating the ratio of correctly segmented nuclei or cells out of the total nuclei in a z-stack. Error bar represents standard deviation across all samples (n=20 growth plates), resulting in on average 50% nuclei and 25% cells correctly segmented from each growth plate sample. (**D**) Nuclei and cells from different growth plate zones were imaged by confocal microscopy, manually segmented, and their volumes and surface areas were compared to those that were segmented semi-automatically from light-sheet images (n=80) using chi-square distance test, resulting in non-significant differences (cell volume, p = 1.00; cell surface area, p = 0.92; nucleus volume, p = 0.22; nucleus surface area, p = 0.24).

**Supplementary Figure 2.**
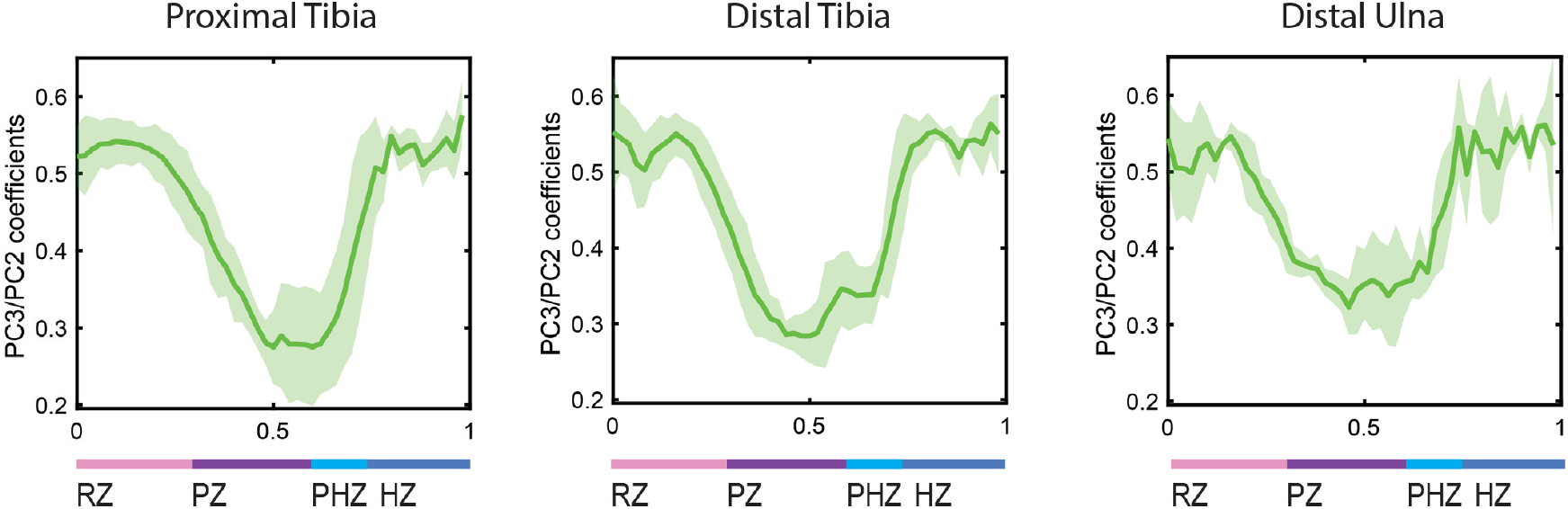
PC coefficient ratios are conserved across different growth plates. The shortest to second longest cell axis ratio (PC3/PC2) was plotted as a spatial profile along the differentiation axis. In all three growth plates, as cells differentiated from the RZ to the PZ, they decreased their PC3/PC2 ratio by half and then returned to the same ratio as the RZ when they differentiated to the HZ. Shaded region shows standard deviation between samples (proximal and distal tibia, n = 5; distal ulna, n = 3).

**Supplementary Figure 3.**
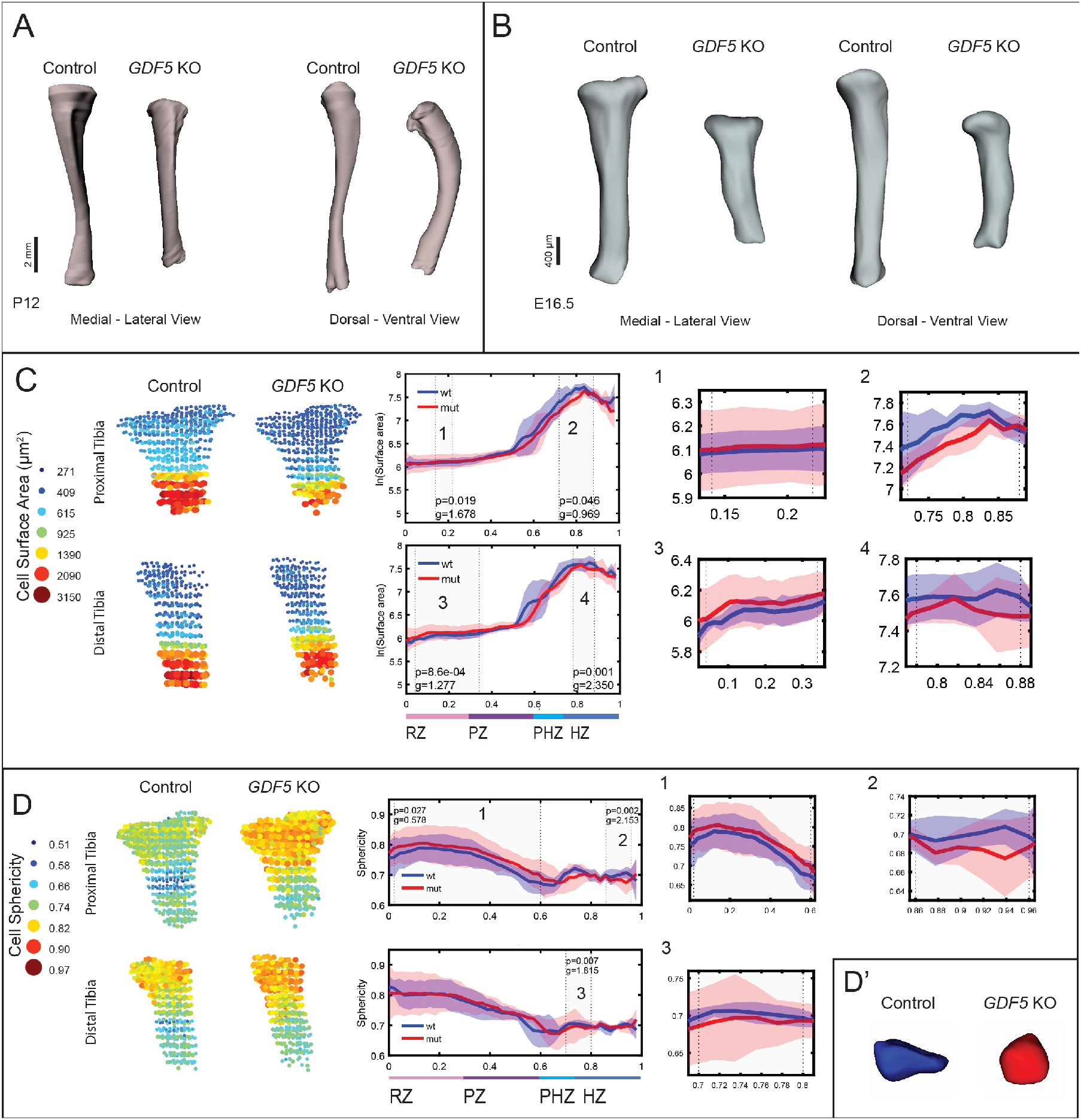
*Gdf5* regulates chondrocyte surface area and sphericity in the growth plate. **(A)** Surface renderings of micro-CT scans of P12 control and *Gdf5* KO tibias show that KO tibias are shorter than controls and have abnormal curvature. **(B)** Surface renderings of light-sheet scans of control and *Gdf5* KO tibias show that the morphological abnormalities are already present at E16.5. (**C** and **D**) Representative 3D maps of control and mutant samples as well as comparative spatial profiles show that *Gdf5* KO tibias display aberrant cell surface area and sphericity. In proximal (PT) and distal (DT) *Gdf5* KO tibial growth plates (**C**), surface area was abnormally high in the RZ (PT p = 1.9e^−02^ g = 1.678; DT p =8.6e^−04^ g = 1.277) and low in the HZ (PT p = 4.6e^−02^ g = 0.969; DT p = 1.0e^−03^ g = 2.35). (**D**) Cells in the RZ and PZ of the *Gdf5* KO PT had significantly higher sphericity (p = 2.7e^−02^ g = 0.578), indicating a loss of polarity and inability to flatten. This is also shown by cell surface renderings from the RZ of control and mutant growth plates (**D’**). Additionally, cells in the HZ of *Gdf5* KO PT and DT growth plates had significantly lower sphericity (PT p = 2.0e^−03^ g = 2.153; DT p = 7.0e^−03^ g = 1.815). In the spatial profiles, wt = control and mut = *Gdf5* KO. p-values were calculated by Student’s *t*-test between mean value of control (n = 5) and mutant samples (n = 4); g, effect size.

## Notes

### Competing Interest Statement

The authors have declared no competing interest.

